# Bird brains fit the bill: morphological diversification and the evolution of avian brain size

**DOI:** 10.1101/2024.07.02.601652

**Authors:** Zitan Song, Szymon M Drobniak, Yang Liu, Carel P van Schaik, Michael Griesser

**Affiliations:** Comparative Socioecology Group, Max Planck Institute for Animal Behavior, Konstanz, Germany; Evolution & Ecology Research Centre, School of Biological, Environmental & Earth Sciences, University of New South Wales, Sydney, Australia; Institute of Environmental Sciences; Jagiellonian University, Krakow, Poland; State Key Laboratory of Biocontrol, Department of Ecology and School of Life Sciences, Sun Yat-sen University, Guangzhou, People’s Republic of China; Department of Evolutionary Anthropology, University of Zurich, Zurich, Switzerland; Centre for Interdisciplinary Study of Language Evolution, University of Zurich, Zürich, Switzerland; Department of Biology, University of Konstanz, Konstanz, Germany; Center for the Advanced Study of Collective Behaviour, University of Konstanz, Konstanz, Germany; Department of Collective Behavior, Max Planck Institute of Animal Behavior, Konstanz, Germany

**Keywords:** sensorimotor complexity and integration, phylogenetic lability analyses, body plan, domain-general intelligence, parental provisioning, social brain, ecological brain

## Abstract

Brain size varies greatly across and even within lineages. Attempts to explain this variation have mostly focused on the role of specific cognitive demands in the social or ecological domain. However, their predictive power is modest, whereas the effects of additional functions, especially sensory information processing and motor control, on brain size remain underexplored. Here, using phylogenetic comparative models, we show that the socio-cognitive and eco-cognitive demands do not have direct links to relative brain size (that is the residual from a regression against body mass) once morphological features are taken into account. Thus, specific cognitive abilities linked to social life or ecology play a much smaller role in brain size evolution than generally assumed. Instead, parental provisioning, generation length, and especially eye size and beak and leg morphology have a strong direct link to relative brain size. Phylogenetic lability analyses suggest that morphological diversification preceded changes in the rate of brain size evolution and greater visual input, and thus that morphological diversification opened up specialized niches where efficient foraging could produce energy surpluses. Increases in brain size provided general behavioural flexibility, which improved survival by reducing interspecific competition and predation, and was made possible by intense parental provisioning. Indeed, comparative analyses in a subset of species show that thicker beaks are associated with larger size of brain regions involved in behavioural flexibility (telencephalon, pallium). Thus, morphological evolution had a key role in niche diversification, which subsequently may have facilitated the evolution of general cognitive flexibility.

## Introduction

Brain size varies greatly across and even within lineages^1^. This fact has attracted ample scientific attention. Among birds and mammals, influential work has focused on the effects of specific eco-cognitive demands and opportunities, including diet^2,3^, food hoarding^4^, extractive foraging^5^, tool use^6^, spatial memory and orientation^7^, as well as socio-cognitive demands, including parenting^8^, social complexity^9,10^ or the establishment and maintenance of social bonds^11^. These effects have been thought to reflect the need for domain-specific cognitive adaptations and were conceptualised in the ecological brain hypothesis^12,13^ or the social brain hypothesis^11^. However, these studies rarely explain a large part of the variation in brain size beyond the effect of body mass^14^. In fact, a recent study on brain size in birds^15^ showed that predictors reflecting eco- and socio-cognitive demands became insignificant when models also included parental provisioning, which estimates the energy supply to the developing brain.

One possible explanation is that the function of brains goes well beyond optimising the balance between traditionally assessed cognitive benefits and the ability to sustain the energetic costs of brains. Brains deal with the perception and processing of sensory information, the cognitive processes involved in planning and decision making, and the control of the resulting motor actions^16^. Accordingly, previous work has revealed strong correlations between brain size and visual input among birds and primates^17,18^ and between brain size and manipulative complexity in primates^19,20^. The previously considered specific eco- and socio-cognitive adaptations may therefore largely be a byproduct of the broader sensorimotor functions of the brain^17,21^. If so, the presumed association between these specific external demands and cognition may be spurious.

Against this background, we here pursue an integrative approach, asking how tightly brain size reflects a species’ overall motor capacities and sensory abilities in birds. We collected data on their brain size, body mass, morphology, life-history, ecology and sociality^15,22^. Birds show large variation in these traits, and their phylogeny is well resolved^23^. Given that birds strongly rely on visual perception^24^, we use eye size as an index of sensory demands^18^. To characterise variation in motor control, we applied Principal Component (PC) analysis to seven morphological variables, reducing data dimensionality in various measures of body plan^25^, for all species for which we had brain size data. We extracted three morphological PCs (mPC) (see Table S1a): species with longer tails, wings and secondary feathers had higher scores on mPC1. Species with wider and deeper, and therefore thicker beaks had higher scores on mPC2. Species with longer beaks and tarsi had higher scores on mPC3. We also applied PC analysis to 18 ecological niche variables, extracting four ecological PCs (ePC): arboreal frugivory (ePC1), aquatic predation (ePC2), land predation (ePC3) and insectivory (ePC4). From the four social niche variables, we extracted two social PCs (sPC): social bonding (sPC1) and gregariousness during breeding and foraging (sPC2) (see Table S1b-c).

We subsequently assessed the statistical effects of these variables and the links among them. We included generation length as a life history proxy^26^, reflecting the positive effect of brain size on survival^27,28^. We also controlled for variables that affect the ability to supply the energy costs of brains^29^, such as duration of parental provisioning^15^ and the presence of long-distance migration^30^. Finally, all analyses controlled for effects of body mass.

## Results

Here we report the results based on the 1155 species for which brain size, social, ecological and morphological variables were available. First, to explore the overall structure of evolutionary correlations among all variables we examined their phenotypic correlations, decomposing them into phylogenetic and non-phylogenetic components of trait associations on the between-species level. We conducted multivariate phylogenetic regressions using PGLS in the package Phylolm, in which each variable was considered a response variable, while all other variables entered into the model as predictors along with log body mass (Fig. 1a). Thus, in those models, each variable was analysed both as a response variable and as a predictor. Figure 1b highlights the significant bivariate relationships (Bonferroni-corrected) depicted in Figure 1a, whereas Figure 1c illustrates the relationships between the key predictors and brain size identified in Figure 1b.

**Figure 1.**
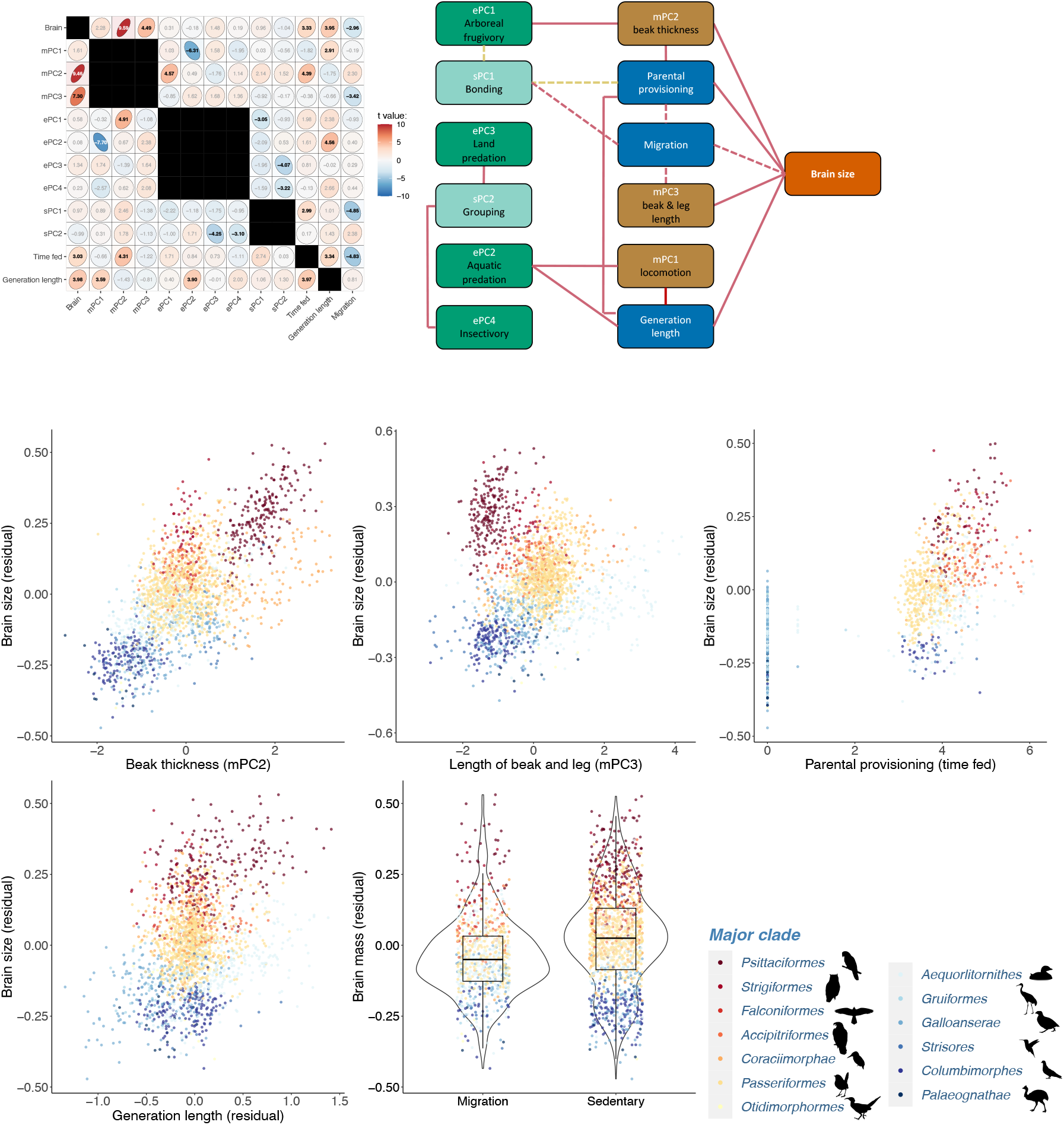
Phylogenetic associations between morphology, life-history traits, eco-social niche and brain size (n =1155). a, presents a heatmap matrix illustrating the relationships between various traits. For each trait in the row, a model was constructed against the other 13 column variables, resulting in a total of 12 models. All models accounted for phylogenetic relationships and included body mass (log-10) as a consistent fixed variable. The models were developed using Phylogenetic Generalized Least Squares (PGLS) via the ‘phylolm’ package. Within each matrix cell, t-values are displayed and their magnitude is visually represented by the color intensity of the cell. Cells with black-colored t-values indicate statistical significance after Bonferroni correction, corresponding to a p-value threshold of less than 0.05/12. b, significant Bonferroni-corrected pair-wise associations among morphology (yellow), life history (blue), ecology and sociality (green) and relative brain size (orange); solid lines represent relationships that are significant in both directions, broken lines represent relationships significant only in one direction (migration could only be included as an independent variable). c, pairwise associations between residual brain size and morphological PC2 (beak depth and width) and PC3 (beak length and leg length), parental provisioning, generation length and migration. Major clades of bird were ordered by average brain size from red to blue.

A striking pattern emerged: brain size underwent tight positive correlated evolution with beak thickness (mPC2) and beak and tarsus length (mPC3), as well as life history variables (increased provisioning time and generation length, and absence of migration), but not with the various eco- or socio-cognitive demands (Fig. 1). Because skull size might constrain brain size^31^, we confirmed that the positive correlation of mPC2 and mPC3 with brain size was not affected by skull size (see Table S2). Fig. 1b also reveals numerous correlations among the categories of variables that confirm known patterns or fit common ecological knowledge, but were not affected by brain size. Concerning the links between morphology and ecology, shorter wings and tails (mPC1) were correlated with aquatic predation (ePC2)^25^ and longer generation length, thicker beaks (mPC2) with arboreal frugivory (ePC1), and longer legs and beaks (mPC3) with migratory lifestyle. Concerning the links between morphology and behaviour, thicker beaks (mPC2) were correlated with longer parental provisioning periods. Concerning the non-morphological variables, preying on terrestrial vertebrates (ePC3) or insects (ePC4) was positively associated with solitary lifestyles (sPC2)^32,33^, the presence of social bonds (sPC1) with parental provisioning and non-migratory lifestyle, and longer generation time with both aquatic predation (ePC2)^34^ and longer parental provisioning. That our analysis recaptures these patterns shows its overall robustness.

We now turn to the evolutionary predictors of brain size. The analyses underlying Figure 1b suggested that any effect of such cognitive demands on brain size acted through morphology or life history. We examined the robustness of this pattern through phylogenetically controlled mixed models in the package MCMCglmm, taking also into account phylogenetic (topological) uncertainty in inter-species relationships. These models estimate the statistical effect on brain size separately for morphological, life-history and eco-social variables, as well as that of all variables combined. The results (Fig. 2, see also Table S3) showed that morphology and life history had stronger correlations with brain size than the socio-ecological predictors. Thus, the combined model explained more variance in relative brain size (R^2^ = 0.52) than the morphological (R^2^ = 0.44), life-history (R^2^ = 0.37), or eco-social model (R^2^ = 0.29). Importantly, the correlations with the latter were no longer significant in the full model. The results remained identical in the smaller dataset (N=660 species) that included eye size (see Fig. S1-S2, Table S4).

**Figure 2.**
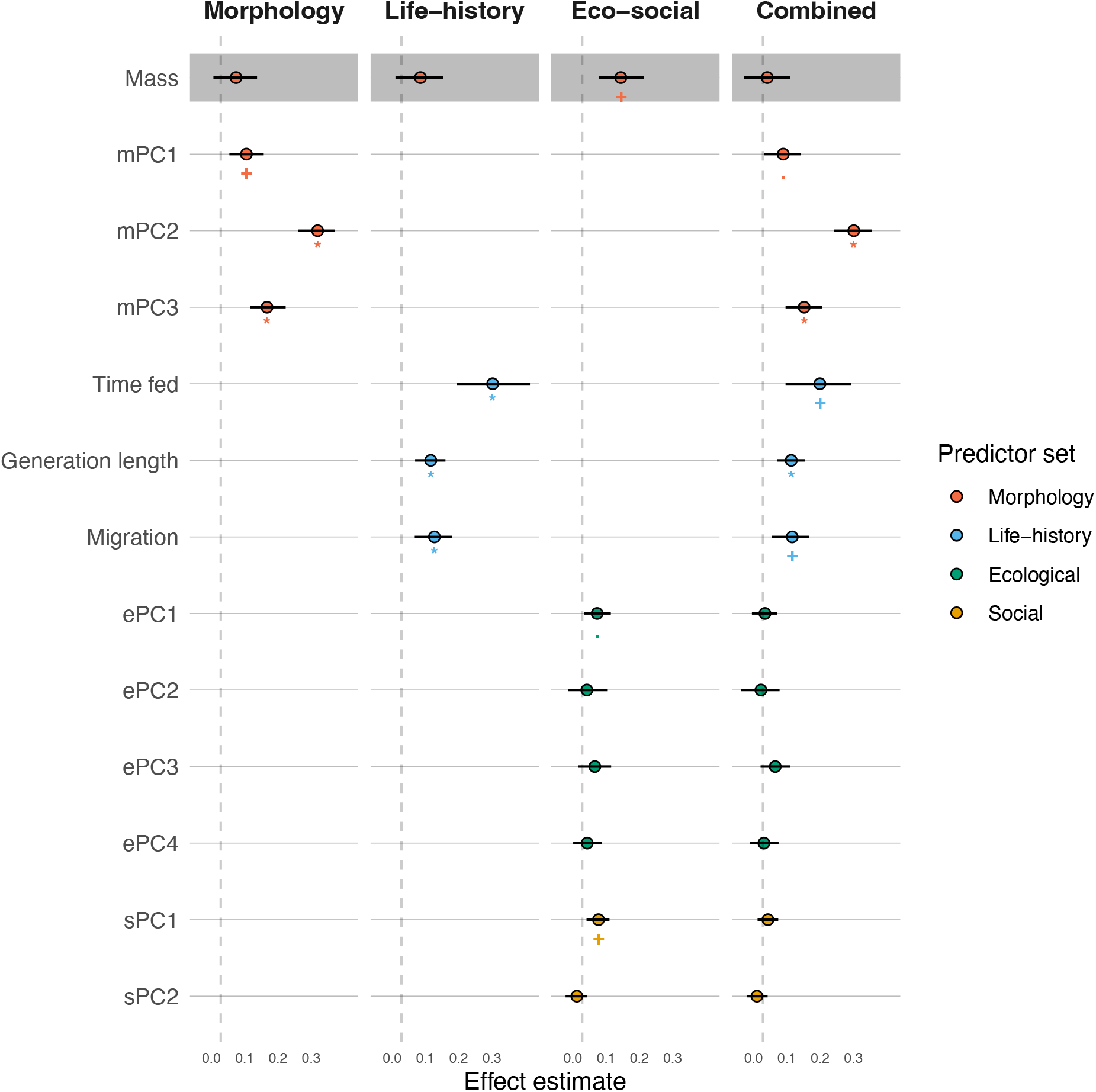
Estimated coefficients and effect sizes of phylogenetically controlled mixed models of the morphology model, the eco-social model, life-history model, and the combined model on relative brain size in birds (n = 1155). Color-filled circles denote estimated effects, lines denote the 95% lower and upper confidence limits generated in the R package MCMCglmm based on a consensus phylogeny. The corresponding full models are shown in Table S4.

We next asked whether the pattern of correlated evolution between brain size and the key predictors of the combined model (Fig. 2) had remained constant throughout evolutionary time. We therefore regressed the ancestral values of each variable against the internal node ages until 57 Myr, and conducted PGLS analyses of the predictors of brain size at one-million years intervals going back in time. The results (Fig. 3) showed that especially beak thickness (mPC2) and parental provisioning period yielded the strongest correlated evolution with brain size throughout the entire period considered. Variables with smaller effect sizes only became significant closer to present time as sample size increased. Thus, in birds, the co-evolution of morphology, life-history and brain size has a deep phylogenetic history.

**Figure 3.**
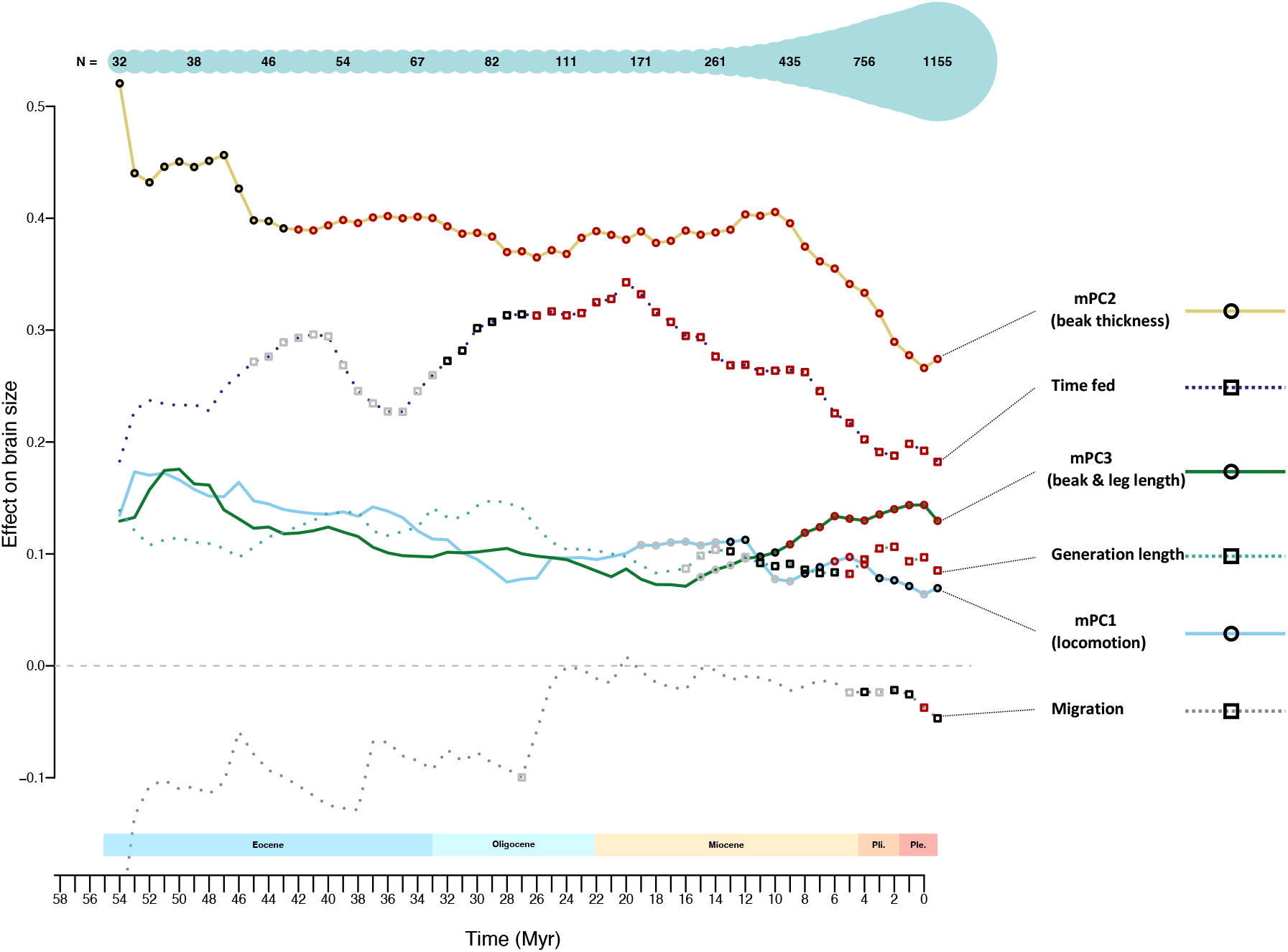
Effect of morphological traits and life-history on relative brain size over evolutionary time. The effects are estimated via pgls models in the ‘phylolm’ package with morphological traits and life-history traits as independent variables and relative brain size as the dependent variable. The significance level is represented by colour: red indicates a p-value < 0.001, black a p-value < 0.01, and grey a p-value < 0.05. Morphological trait values in deep time were determined via ancestral state reconstruction in the package ‘phytools’ using a variable-rates model, with calculations performed at 1 million-year intervals. A time-calibrated maximum-likelihood phylogenetic tree was pruned based on the time when ancestral states were estimated. Note that the analysis starts at 57 Ma as before then as the sample size was too low before (N < 30).

So far, our analyses only considered correlated evolution and therefore could not assign selective agency to any of the variables involved. We therefore wished to identify variables whose rate of evolutionary change shifted significantly before similar changes in other correlated characters. We hypothesised that the former variables would have driven the changes in the latter. Accordingly, we conducted phylogenetic analyses that reconstructed the rate of change per unit time of relative brain size and key predictors over evolutionary time, using the BAMM. These analyses (Fig. 4) show that morphology, especially the length of legs and beaks (mPC3) diversified particularly during the Paleocene and Eocene. After that, the overall morphological diversification continued, whereas change in brain size began to catch up during the Oligocene, suggesting it responded to the newly modified body plans (Fig. 4c). Subsequently the response gradually turned into more tightly correlated evolution between brain size and morphology, suggesting increased integration between morphology and brain functions. When we added eye size to the set of variables, its rate of change was very similar to that of the brain (Fig. S3), suggesting that the processing of sensory inputs is a major component driving the evolutionary response of brain size.

**Figure 4.**
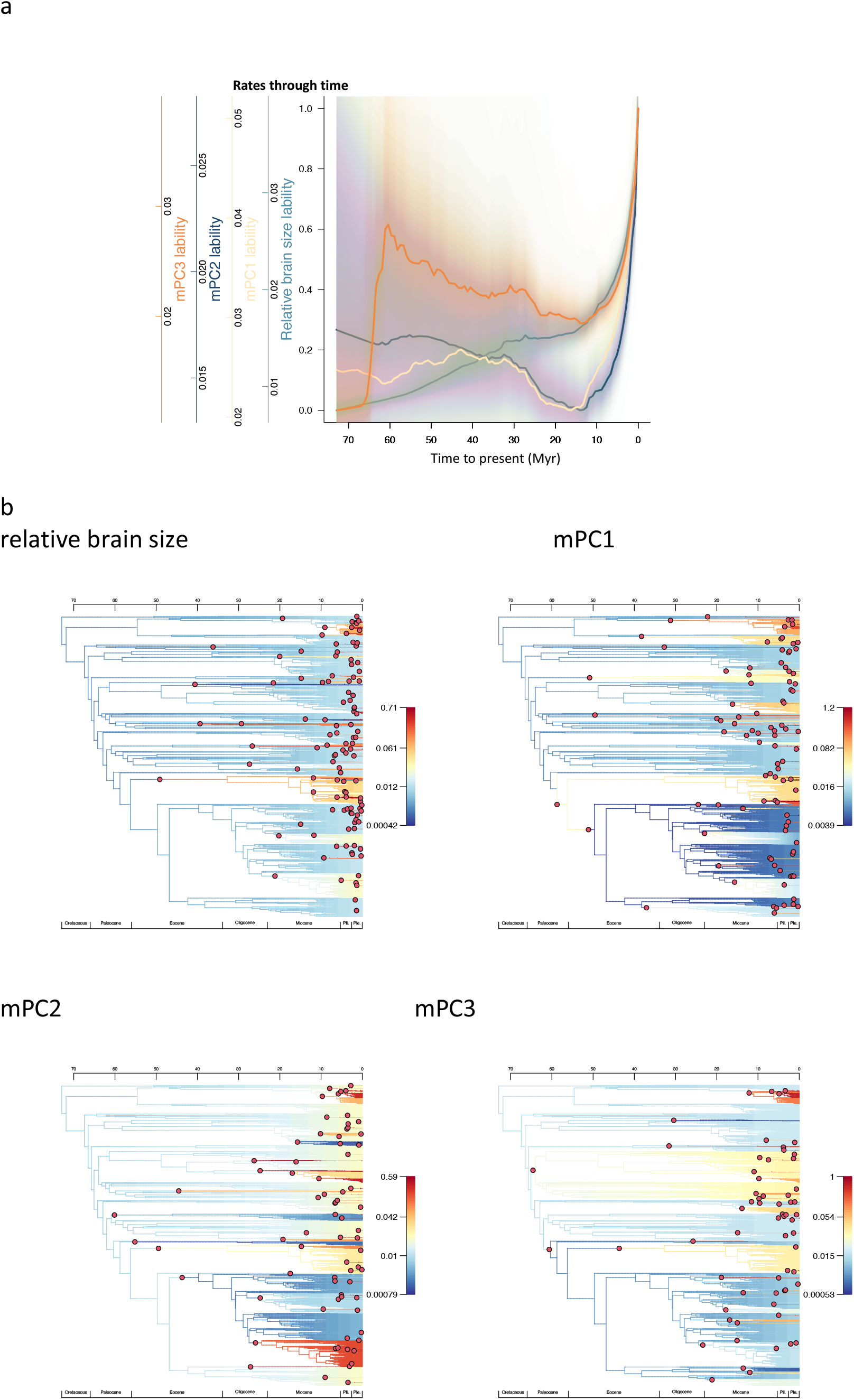
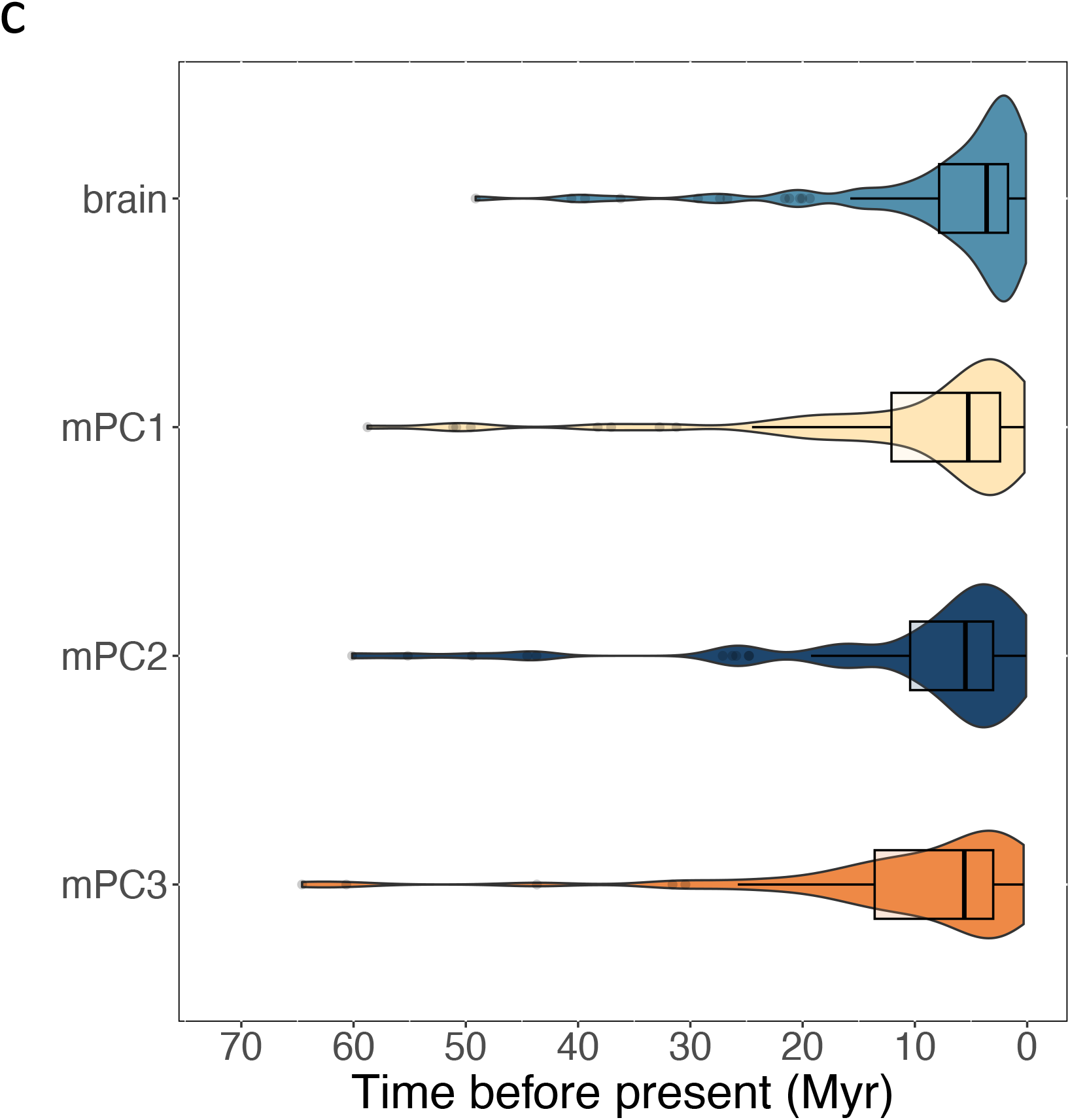
a) median rates and rate distributions for net phenotypic macroevolutionary rate. the y axis is scaled from zero to the maximum median rate for all dataset, for relative comparability. b) Rates of phenotypic (brain and mPC1-3) evolution in bird, branches are colored in a rainbow scale by Jenks natural breaks method. Red dots at nodes represent diversification shifts in the maximum credibility set. c) box plot showing the distribution of major shifts in best posterior probability shift configuration, the major shifts of mPCs is earlier than brain (Mann-Whitney U test: mPC3: W = 2783, p =0.017; mPC2: W = 3426, p =0.037; mPC1: W = 4500, p =0.045).

The above detailed results suggest a key role for beak morphology and sensory input for brain size evolution. This pattern may reflect that thicker beaks require increased fine-motoric abilities, and thus a larger cerebellum. Alternatively, thicker beaks provide the greater morphological divergence from potential competitors via the refined decision making needed to thrive in secure niches, and thus require a larger telencephalon and pallium. Additional analyses in 110 species where detailed data on brain regions were available showed that ticker beaks (mPC2) are associated with both larger relative telencephalon and pallium mass, as well as neuron numbers (Fig. 5, Table S5), whereas longer wings and tails (mPC1) were associated with larger relative cerebellum mass and neuron numbers. Thus, these patterns support the notion that morphology provides the foundation for the niche whereas general behavioural flexibility is responsible for the performance in this niche.

**Figure 5.**
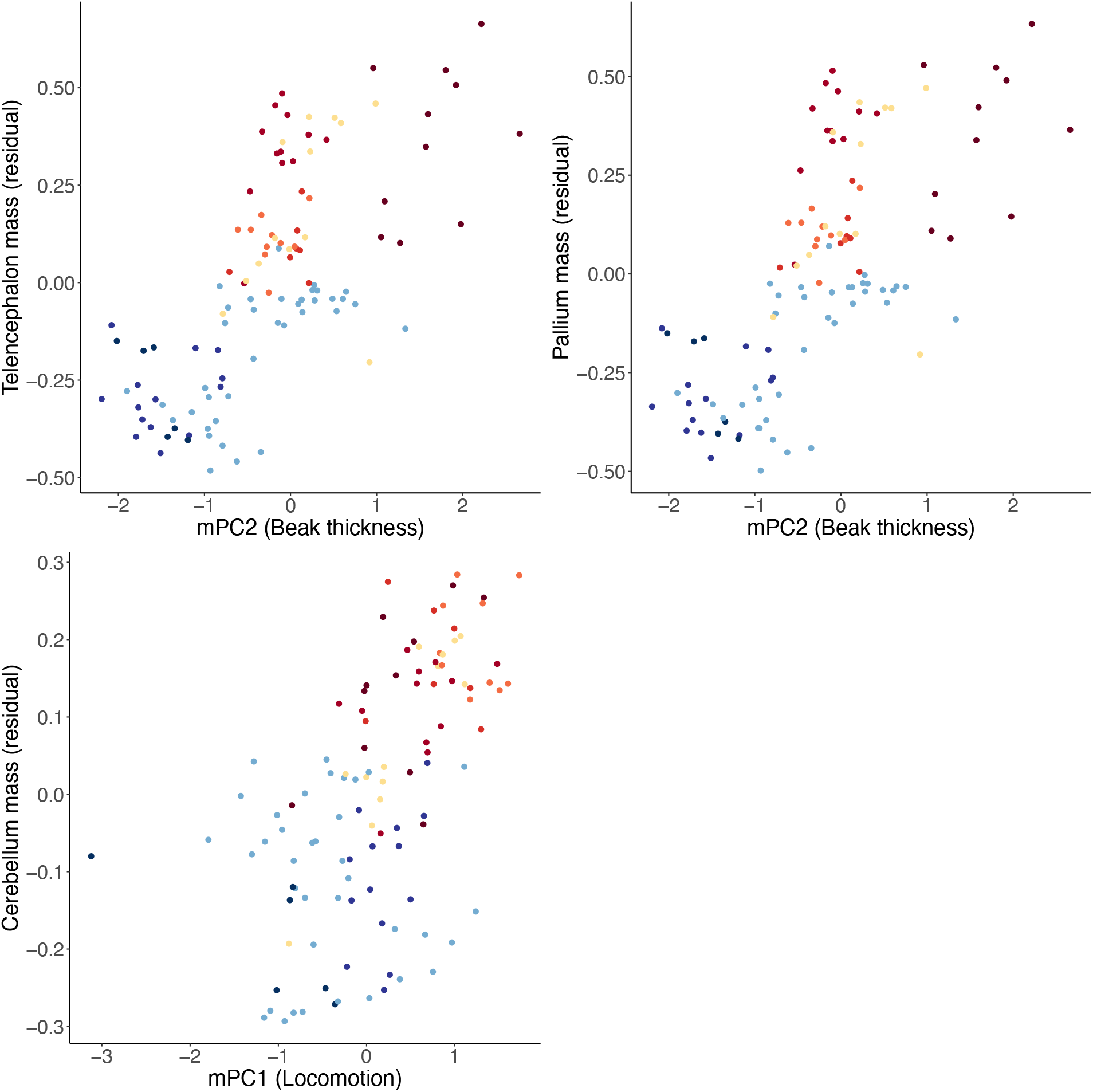
Pairwise associations between residual brain region size and morphological PC2 (beak depth and width) and PC1 (locomotion). Major clades of bird were ordered by average brain size from red to blue similar as in Figure 1c.

## Discussion

Our analysis of brain size variation in birds revealed that body plan measures (morphology) and the amount of sensory inputs (here indexed by eye size) showed the strongest correlated evolution with brain size (Fig. 1-2, S1-S2). In contrast, previously considered socio- and eco-cognitive challenges did not have any direct links to brain size when sensorimotor measures were also included in the model. Independently thereof, energetic aspects remained important as well^27^: migratory species had smaller brains than sedentary species, whereas those with larger brains engaged in extended parental provisioning (Fig. 2).

We found that evolutionary changes in the rate of morphological evolution preceded changes in the rate of evolution of brain size and eye size (Fig.4&S3), suggesting that particular morphologies facilitated the evolution of greater visual sensory and processing abilities, allowing animals to perceive the world in finer detail (Fig. S1-S2). Simultaneously, adaptive changes in brain size and composition (Fig. 5) then enabled individuals to effectively process this information, make adequate action decisions and execute the proper motor actions to succeed in their niche. We conclude that sensing, information processing, and acting became tightly integrated through coevolution, but the triggering changes were morphological.

The important causal role of body plan in evolutionary changes in the niche and brain size is reflected in the phylogenetic history of birds. Ancestral bird lineages were predominantly precocial and terrestrial, with simple foraging niches and short, flat beaks used for picking up food items. Ducks expanded their niches to aquatic habitats, which did not require much morphological change^25^. All of these species have relatively low levels of parental provisioning and small brains^15^. The advent of Neoaves in the early Cenozoic 65 Myr ago^23^ led to a diversification of beak morphology, consistent with the increase in overall beak volume, and the associated foraging niches^25^, including extended flight as well as arboreal nesting and foraging. These new niches supported the consumption of novel foods, while the security of arboreal nesting enabled the evolution of both slower life-histories^35^ and post-hatching provisioning^15,36^, together facilitating brain size evolution. Thus, this pattern indicates a gradual increase in integration between sensorimotor capacities and morphology. This niche construction perspective^37^ on brain size evolution suggests the value of shifting the focus from meeting specific socio- and eco-cognitive challenges towards a more holistic perspective on functioning in the niche, with respect to energy balance and survival.

The minor and at best indirect role in brain size evolution of specific social or ecological cognitive demands relative to that of body plan and sensory inputs may be explained as follows. The fundamental control of running a particular body plan may already cover all niche-related cognitive demands because the above-mentioned planning and adjustment requires processes normally encompassed under the rubric of cognition^16^. Thus, increasingly fine adjustment to a specific niche can be reached through domain-general cognitive abilities ^38,39^, which turn innate preferences for actions or contexts into skills through learning from caregivers during development^40,41^.

Our findings therefore contrast with previous work that attributed a major role in brain size evolution to specific socio-or eco-cognitive challenges (e.g., social bonding, social complexity, diet, extractive foraging, habitat complexity) in specific lineages (e.g., primates, carnivores, corvids or parrots^3,42,43^. This approach is less successful when generalised to other lineages. For example, while social challenges appear to affect brain size in primates^43^, they do not appear to do so in carnivores^42,44^. Likewise, diet was found to affect brain size in primates^3^ but not in marsupials^45^. The present results suggest that the relative homogeneity of body plan and sensory abilities within a single taxon may have masked their influences on brain size and allowed the modest effects of cognitive challenges to be expressed. We therefore conclude that the role of sensorimotor complexity and integration in brain size evolution is almost certainly far greater than that of the often highlighted specific cognitive adaptations underlying features such as social bonding, tool use or diet switching.

## Methods

### Data collection

Data on brain size were collected from two published sources for 2135 bird species^15,22^. For all 2135 bird species, we selected 7 key morphological variables from AVONET^46^: wing length, secondary feather length, tail length, tarsus length, and the width, length to culmen and depth of the beak. We also obtained standardised scores for the ecological niche of each species by using published data on diet categories (N=10) and foraging stratum (N=7)^47^. Since the degree of food processing complexity can vary based on ecology and foraging niche, which may also affect brain size, we also included food handling levels as a variable that measures the number of processing steps required to extract food^15,48^.

We collected social niche variables from published sources^15,49^: social bond strength, number of parental caretakers, sociality during the non-breeding and the breeding season. Social bond strength distinguished species with short bonds during mating only, seasonal bonds, and long-term bonds that extend beyond a single breeding season. As to the mean number of caretakers, mound-nesting species had zero caretakers, species with uniparental care one, species with bi-parental care two, and cooperatively breeding species the mean value of caretakers. For the sociality during the non-breeding season, we separated asocial species, pair living species, species that live in small groups (usually 3 to 30 individuals; personalized groups where group members know each other, e.g., family groups), and large groups (usually more than 30 individuals; ephemeral and or anonymous groups^50^. Sociality during the breeding season distinguished among species that breed singly and those that breed in colonies.

The parental provisioning period included both the time young are fed inside and outside the nest^15^. Imputed generation lengths of all species were collected from^26^. Data on migratory habits were collected from^15^, differentiating between sedentary species (including local movements) and migratory species (short- and long-distance migrants, altitudinal migrants). We also gathered data on the axial length (mm) of bird eyes from multiple sources (see supplementary data for details). For those species where multiple measurements of eye dimensions were available in the literature, we calculated the mean values of all provided measurements.

Cranial morphological traits (beak depth and width) are highly correlated with skull size, which in turn is positively correlated with brain size. To check for a possible constraining effect of skull size, we gathered data on the height, width and length of the cranium for N=667 species from https://skullsite.com. Table S2 shows that skull size did not affect the relationship between the morphological principal components and brain size. The size and the number of neurons in the telencephalon, pallium, cerebellum and brainstem (N=110 species) were collected from a published dataset^51^.

We note that the sample sizes for the different variables varied due to variable data availability (see in the tables and figures for the respective sample size).

### Data analysis

All statistical analyses were carried out in the R 4.1.1 environment^52^. Individual morphological, ecological and social parameters are usually correlated to other predictors of the same category, and thus, we applied a Principal Component Analysis (PCA) approach to reduce the dimensionality of these datasets. Given the allometric relationship of body mass with other morphological predictors, we calculated their residuals from regressions against body mass after log-10 transformation. PCAs were calculated with the package psych^53^ using varimax rotation. For morphology, we extracted 3 PCs: (locomotion (mPC1), beak thickness (mPC2), beak and leg length (mPC3) (Table S1a). For diet and foraging strata, we extracted 4 PCs: arboreal frugivory (ePC1) aquatic predation species (ePC2), land predation (ePC3), and insectivory (ePC4) (Table S1b). For social parameters, we extracted 2 PCs: social bonding (sPC1), and social breeding (sPC2) (Table S1c). Generation length, skull size and eye size were all correlated with species body mass, and thus we took their residuals from regressions against body mass (log-10 transformed). We excluded the two kiwi species from all subsequent analyses because they virtually lack wings, making them extreme outliers in the mPC1. Including the two species prevented the convergence of ancestral state reconstruction (see below for details).

### Bivariate phylogenetic relationships among factors

We fitted 12 phylogenetic generalised least squares (PGLS) models in the package phylolm^54^ to investigate the bivariate correlations between brain size, morphological predictors, eco-social factors, and life-history traits (Table S6, all traits were scaled and centred before the analyses). In each model, a different variable was designated as the response, while the remaining variables were included as predictors. In all models, log-transformed body mass was included as a fixed variable to control for the nonlinear effects of allometry. We did not consider the relationships among morphology traits, ecology niches, and social niches, as our interest lies in the relationships between morphology, life-history traits, eco-social niche, and brain size, rather than within these categories. We also included eye size as an indicator of sensory input and repeated the above detailed procedure to assess its associations with morphology, life-history traits, eco-social niche and brain size (N=660 species, Table S7).

We controlled for phylogeny using an well-resolved avian phylogenetic tree^23^, matching the tips to the phylogeny of Jetz etal.^55^, following the methodology detailed in Cooney et al.^56^.

### Test the effects of predictors on brain

We fitted 4 phylogenetically controlled mixed models (N=1155 species) that assessed the dependence of relative brain size on morphology predictors (model 1), eco-social predictors (model 2) life-history predictors (model 3), and of all predictors (model 4; see S3-4 Tables for a full list of predictors in each of the models). All models were fitted using the package MCMCglmm^57^. We used chains of 50,000 iterations, with the first 10,000 iterations discarded as burn-ins and thinned every 40 iterations. Inspection of the final MCMC samples did not show any sign of autocorrelation. All effective sample sizes were close to the actual numbers of sampled draws from predictors’ posteriors (S3-4 Table). In all models, inverse-gamma priors were used for residual variances (parametrized as inverse-Wishart with V = 1 and ν = 0.002). The prior for phylogenetic effect was formed as a weakly informative half-Cauchy density (parameter expanded priors with V = 1, ν = 1, α_μ_ = 0 and αV = 10,000). Priors for fixed effects were left as default (Gaussian densities with μ = 0 and large variance). We used a modification of the variance inflation factor (VIF) analysis adjusted to phylogenetic comparative mixed models (https://github.com/mrhelmus/phylogeny_manipulation), finding a low degree of collinearity (all VIF values <2; S8 Table).

We fitted a phylogenetically controlled mixed model to test whether the effects of morphological predictors on brain size covary with skull size. This model was implemented in MCMCglmm (N = 667) with residual brain size as the response variable and body mass, mPC1-3, and residual skull size as the dependent variables. We also fitted MCMCglmm models (N = 110) to test the effects of morphological predictors on the size and number of neurons in different brain regions, including the telencephalon, pallium, cerebellum, and brainstem. Residuals for all brain regions were calculated against body mass. Since the effects of morphology on brain regions may covary with parental provision, we considered the developmental mode (precocial vs. altricial) as a dependent variable.

### Effects of predictors on brain size through time

The effects of predictors on brain size may change over evolutionary time. Thus, we reconstructed ancestral states of key traits and assessed their effects of predictors on brain size through time. We first reconstructed the ancestral-state of key traits that the above analyses (Table S3/Figure 1&2) showed to have direct effects on brain size (mPC1-3, parental provision time, generation length, migration, N = 1155) following the approach used by Cooney et al.^56^. For each trait, we assessed the phylogenetic signal by calculating *Pagel’s lambda* and *Blomberg’s K* in the package phytools *v. 1*.*0*.*1*^58^. We then tested the fit of 4 evolutionary models of trait evolution for each of the four traits: a white noise, a Brownian motion, a single-optimum Ornstein–Uhlenbeck and early burst of trait evolution, using the function fitContinuous of the package geiger *v. 2*.*0*.*10*^59^. Additionally, we fitted a Brownian motion model, which allows for rate shift on branches and nodes using the software BayesTrait *v. 3* (http://www.evolution.rdg.ac.uk/)^60^ with uniform prior distributions adjusted to our dataset and applying single-chain Markov-chain Monte Carlo runs with at least 5 x 10^8^ iterations (detail see Table S9). We sampled parameters every 10,000th iteration, after a burn-in period of at least 10,000,00 iterations. For all continuous traits, we applied independent contrast models, for migration (a binary trait), we used a multistate model. We then tested for each trait for convergence of the chain using the Cramer–von Mises statistic implemented in the package coda *v. 0*.*19-4*^61^. The models were compared by calculating their log-likelihood and Akaike information criterion (AIC) difference (Table S10). Based on differences in AIC, the variable-rates model was best supported for all traits (Table S10).

We reconstructed ancestral states for each node of key traits using the mean rate-transformed tree derived from the variable-rates model. We then slice the original species tree with 1Myr time intervals and calculated the ancestral states at each time point.

For each time point we extracted the branches existing at that time and predicted the trait value via linear interpolation between nodes. We subsequently assessed the effect of morphological, ecological, social, and life-history variables on brain size. To achieve this, we constructed Phylogenetic Generalized Least Squares (PGLS) models for each 1Myr time interval using the ‘phylolm’ package^54^. Since the number of reconstructed ancestral species increases over time, we only conducted the PGLS analysis after 57Myr years, when the sample size exceeded 30. The ancestral states for brain size as well as all other traits were scaled and centred for the analysis.

### Macroevolutionary Rate Analysis

We used the package BAMMtools^62^ along with a set of parametric and semiparametric test statistics to analyse macroevolutionary rates of brain size and key traits related to it, using N=2133 species after two Kiwis been removed. All traits were scaled and centred before the analyses to facilitate comparison of rate shifts among them. We used the BAMM trait model to reconstruct evolutionary rates through time for residual brain size and mPC1-3. We anticipated a total of 20 shifts and configured the priors settings using the package BAMMtools^62^ via the function setBAMMpriors. We ran 10^9^ generations of Markov chain Monte Carlo (MCMC) in four chains with δT set as 0.1 and the swap period as 10,000 generations, sampling every 10,000th generation. MCMC scaling, move frequency, and initial values of parameters followed recommendations in BAMM documentation. The first 10% of generations were discarded as burn-in. To render these rates comparable, we scaled them to a proportion of the maximum (that is contemporary) rates. The effect sizes of the number of events and log likelihoods are presented in Table S11, all exceeding 1000. We processed the BAMM output files with BAMMtools^62^, applying a 10% burn-in.

We calculated the macroevolutionary rates of traits over time using the *getRateThroughTimeMatrix* function from BAMMtools^62^ and plotted the rates of all traits, scaling them from zero to the highest median rate to compare the evolutionary dynamics between brain size and morphology. In addition, we identified the nodes most likely to represent significant rate shifts using the *getBestShiftConfiguration* function in BAMMtools^62^. We then compared the timing of major rate shifts between morphology and brain size using a Wilcoxon test. We also performed the above described BAMM analyses using a smaller dataset (N=1199 species) that included eye size (residual), in conjunction with brain size and morphology. We anticipated a total of 10 shifts, while the other settings were the same as those above.

## References

1. Jerison, H. Evolution of the Brain and Intelligence. (Academic Press, New York, 1973).

2. Harvey, P. H., Clutton-Brock, T. H. & Mace, G. M. Brain size and ecology in small mammals and primates. Proc Natl Acad Sci U S A 77, 4387–4389 (1980).

3. DeCasien, A. R., Williams, S. A. & Higham, J. P. Primate brain size is predicted by diet but not sociality. Nat Ecol Evol 1, 0112 (2017).

4. Garamszegi, L. Z. & Eens, M. The evolution of hippocampus volume and brain size in relation to food hoarding in birds. Ecol Lett 7, 1216–1224 (2004).

5. Parker, S. T. & Gibson, K. R. Object manipulation, tool use and sensorimotor intelligence as feeding adaptations in cebus monkeys and great apes. J Hum Evol 6, 623–641 (1977).

6. Lefebvre, L., Nicolakakis, N. & Boire, D. Tools and brains in birds. Behaviour 139, 939–973 (2002).

7. Sherry, D. F., Jacobs, L. F. & Gaulin, S. J. C. Spatial memory and adaptive specialization of the hippocampus. Trends Neurosci 15, 298–303 (1992).

8. Gittleman, J. L. Female brain size and parental care in carnivores. Proc Natl Acad Sci U S A 91, 5495–5497 (1994).

9. Burish, M. J., Kueh, H. Y. & Wang, S. S.-H. Brain Architecture and Social Complexity in Modern and Ancient Birds. Brain Behav Evol 63, 107–124 (2004).

10. Ashton, B. J., Thornton, A. & Ridley, A. R. An intraspecific appraisal of the social intelligence hypothesis. Philos Trans R Soc Lond B Biol Sci 373, 20170288 (2018).

11. Dunbar, R. I. M. The social brain hypothesis. Evol Anthropol 6, 178–190 (1998).

12. Milton, K. Distribution patterns of tropical plant foods as an evolutionary stimulus to primate mental development. Am Anthropol 83, 534–548 (1981).

13. Byrne, R. W. The Technical Intelligence hypothesis: An additional evolutionary stimulus to intelligence? in Machiavellian Intelligence II 289–311 (Cambridge University Press, 1997).

14. Wartel, A., Lindenfors, P. & Lind, J. Whatever you want: Inconsistent results are the rule, not the exception, in the study of primate brain evolution. PLoS One 14, e0218655 (2019).

15. Griesser, M., Drobniak, S. M., Graber, S. M. & van Schaik, C. Parental provisioning drives brain size in birds. Proc Natl Acad Sci U S A 120, e2121467120 (2023).

16. Shettleworth, S. J. Cognition, Evolution, and Behavior. (Oxford University press, Oxford, 2009).

17. Kirk, E. C. Visual influences on primate encephalization. J Hum Evol 51, 76–90 (2006).

18. Garamszegi, L. Z., Møller, A. P. & Erritzøe, J. Coevolving avian eye size and brain size in relation to prey capture and nocturnality. Proc R Soc Lond B Biol Sci 269, 961–967 (2002).

19. Heldstab, S. A., Isler, K., Schuppli, C. & van Schaik, C. P. When ontogeny recapitulates phylogeny: Fixed neurodevelopmental sequence of manipulative skills among primates. Sci Adv 6, (2020).

20. Heldstab, S. A. et al. Manipulation complexity in primates coevolved with brain size and terrestriality. Sci Rep 6, 1–9 (2016).

21. Catania, K. C. & Kaas, J. H. The unusual nose and brain of the star-nosed mole. Bioscience 46, 578–586 (1996).

22. Sayol, F., Downing, P. A., Iwaniuk, A. N., Maspons, J. & Sol, D. Predictable evolution towards larger brains in birds colonizing oceanic islands. Nat Commun 9, 2820 (2018).

23. Prum, R. O. et al. A comprehensive phylogeny of birds (Aves) using targeted next-generation DNA sequencing. Nature 526, 569–573 (2015).

24. Martin, G. R. The Sensory Ecology of Birds. (Oxford University Press, Oxford, 2017).

25. Pigot, A. L. et al. Macroevolutionary convergence connects morphological form to ecological function in birds. Nat Ecol Evol 4, 230–239 (2020).

26. Bird, J. P. et al. Generation lengths of the world’s birds and their implications for extinction risk. Conservation Biology 34, 1252–1261 (2020).

27. Isler, K. & van Schaik, C. P. The expensive brain: a framework for explaining evolutionary changes in brain size. J Hum Evol 57, 392–400 (2009).

28. Minias, P. & Podlaszczuk, P. Longevity is associated with relative brain size in birds. Ecol Evol 7, 3558–3566 (2017).

29. Heldstab, S. A., Isler, K., Graber, S. M., Schuppli, C. & van Schaik, C. P. The economics of brain size evolution in vertebrates. Curr Biol 32, R697–R708 (2022).

30. Vincze, O. Light enough to travel or wise enough to stay? Brain size evolution and migratory behavior in birds. Evolution 70, 2123–2133 (2016).

31. Holekamp, K. E., Swanson, E. M. & Van Meter, P. E. Developmental constraints on behavioural flexibility. Philos Trans R Soc Lond B Biol Sci 368, 20120350 (2013).

32. Song, Z., Liker, A., Liu, Y. & Székely, T. Evolution of Social Organization: Phylogenetic Analyses of Ecology and Sexual Selection in Weavers. Am Nat 200, 250–263 (2022).

33. Beauchamp, G. Flocking in birds is associated with diet, foraging substrate, timing of activity, and life history. Behav Ecol Sociobiol 76, 74 (2022).

34. Hamer, K. C. & Schreiber, E. A. Breeding biology, life histories, and life history-environment interactions in seabirds. in Biology of marine birds (eds. Schreiber, E. A. & Burger, J.) 217–261 (CRC Press, 2001).

35. Pomeroy, D. Why fly? The possible benefits for lower mortality. Biological Journal of the Linnean Society 40, 53–65 (1990).

36. van Schaik, C. P. et al. Extended parental provisioning and variation in vertebrate brain sizes. PLoS Biol 21, e3002016 (2023).

37. Odling-Smee, J., Erwin, D. H., Palkovacs, E. P., Feldman, M. W. & Laland, K. N. Niche Construction Theory: A Practical Guide for Ecologists. Q Rev Biol 88, 3–28 (2013).

38. Burkart, J. M., Schubiger, M. N. & van Schaik, C. P. The evolution of general intelligence. Behav Brain Sci 40, e195 (2017).

39. Ashton, B. J., Ridley, A. R., Edwards, E. K. & Thornton, A. Cognitive performance is linked to group size and affects fitness in Australian magpies. Nature 554, 364–367 (2018).

40. Uomini, N., Fairlie, J., Gray, R. D. & Griesser, M. Extended parenting and the evolution of cognition. Philos Trans R Soc Lond B Biol Sci (2020).

41. Slagsvold, T. & Wiebe, K. L. Learning the ecological niche. Proc R Soc Lond B Biol Sci 274, 19–23 (2007).

42. Finarelli, J. A. & Flynn, J. J. Brain-size evolution and sociality in Carnivora. Proc Natl Acad Sci U S A 106, 9345–9349 (2009).

43. Shultz, S. & Dunbar, R. I. M. Social bonds in birds are associated with brain size and contingent on the correlated evolution of life-history and increased parental investment. Biol J Linn Soc Lond 100, 111–123 (2010).

44. Holekamp, K. E. Questioning the social intelligence hypothesis. Trends Cogn Sci 11, 65–69 (2007).

45. Todorov, O. S. et al. Testing hypotheses of marsupial brain size variation using phylogenetic multiple imputations and a Bayesian comparative framework. Proc R Soc Lond B Biol Sci 288, rspb.2021.0394 (2021).

46. Tobias, J. A. et al. AVONET: morphological, ecological and geographical data for all birds. Ecol Lett 25, 581–597 (2022).

47. Wilman, H. et al. EltonTraits 1.0: Species-level foraging attributes of the world’s birds and mammals. Ecology 95, 2027–2027 (2014).

48. Schuppli, C., Isler, K. & van Schaik, C. P. How to explain the unusually late age at skill competence among humans. J Hum Evol 63, 843–850 (2012).

49. Tobias, J. A. et al. Territoriality, Social Bonds, and the Evolution of Communal Signaling in Birds. Front Ecol Evol 4, (2016).

50. Griesser, M., Drobniak, S. M., Nakagawa, S. & Botero, C. A. Family living sets the stage for cooperative breeding and ecological resilience in birds. PLoS Biol 15, e2000483 (2017).

51. Sol, D. et al. Neuron numbers link innovativeness with both absolute and relative brain size in birds. Nat Ecol Evol 6, 1381–1389 (2022).

52. Team, R. C. R: A language and environment for statistical computing. (2013).

53. Revelle, W. psych: Procedures for personality and psychological research. Preprint at (2017).

54. Ho, L. si T. & Ane, C. A Linear-Time Algorithm for Gaussian and Non-Gaussian Trait Evolution Models. Syst Biol 63, 397–408 (2014).

55. Jetz, W., Thomas, G. H., Joy, J. B., Hartmann, K. & Mooers, A. O. The global diversity of birds in space and time. Nature 491, 444–448 (2012).

56. Cooney, C. R. et al. Mega-evolutionary dynamics of the adaptive radiation of birds. Nature 542, 344–347 (2017).

57. Hadfield, J. D. MCMC methods for multi-response generalized linear mixed models: the MCMCglmm R package. J Stat Softw 33, 1–22 (2010).

58. Revell, L. J. phytools: an R package for phylogenetic comparative biology (and other things). Methods Ecol Evol 3, 217–223 (2012).

59. Harmon, L. J., Weir, J. T., Brock, C. D., Glor, R. E. & Challenger, W. GEIGER: investigating evolutionary radiations. Bioinformatics 24, 129–131 (2008).

60. Pagel, M. & Meade, A. Bayesian Analysis of Correlated Evolution of Discrete Characters by Reversible-Jump Markov Chain Monte Carlo. Am Nat 167, 808–825 (2006).

61. Plummer, M., Best, N., Cowles, K. & Vines, K. CODA: convergence diagnosis and output analysis for MCMC. R New 6, 7–11 (2005).

62. Rabosky, D. L. et al. BAMMtools: an R package for the analysis of evolutionary dynamics on phylogenetic trees. Methods Ecol Evol 5, 701–707 (2014).

