## Supplemental Table 1-11; Supplemental figure 1-3 for "Bird brains fit the bill: morphological diversification and the evolution of avian brain size"

### Supplementary materials

#### Tables:

**Table S1.** Principal Component Analysis (PCA) loadings and variance explained of morphological traits (A), ecological traits (B), and social traits (C), using the principal function in the package psych[1] with varimax rotation. We applied Standardized loadings of the key contributors to each PC are highlighted in bold. Both kiwi species were removed before the PCA, given their short wing lengths were extreme outliers.

| A) Morphological traits (residual) | **mPC1: locomotion** | **mPC2: beak thickness** | **mPC3: beak & leg length** |
|---|---|---|---|
| **Secondary feather length** | **0.92** | 0.08 | 0.19 |
| **Wing length** | **0.88** | 0.06 | 0.01 |
| **Tail length** | **0.86** | 0.18 | -0.05 |
| **Beak width** | 0.14 | **0.92** | 0.01 |
| **Beak length** | -0.07 | 0.40 | **0.77** |
| **Beak depth** | 0.16 | **0.92** | 0.02 |
| **Tarsus length** | 0.19 | -0.40 | **0.76** |
| *SS loadings* | *2.45* | *2.05* | *1.21* |
| Proportion Variance | *0.35* | *0.29* | *0.17* |
| *Cumulative* Variance | *0.35* | *0.64* | *0.82* |
| *Proportion Explained* | *0.43* | *0.36* | *0.21* |
| *Cumulative Proportion* | *0.43* | *0.79* | *1.00* |

| B) Ecology | **ePC1: arboreal frugivory** | **ePC2: aquatic predation** | **ePC3: land predation** | **ePC4: insectivory** |
|---|---|---|---|---|
| diet: | | | | |
| **Invertebrates** | -0.37 | -0.10 | -0.18 | **0.79** |
| **Mammals, birds** | 0.02 | -0.09 | **0.72** | 0.00 |
| **Reptiles, amphibians** | -0.03 | 0.03 | **0.60** | 0.12 |
| **Fishes** | -0.03 | **0.89** | 0.11 | 0.06 |
| **Vertebrates in general** | -0.01 | -0.01 | 0.08 | -0.02 |
| **Scavenging** | -0.10 | -0.06 | -0.07 | -0.13 |
| **Fruits** | **0.74** | -0.16 | 0.00 | -0.25 |
| **Nectar** | 0.23 | -0.14 | -0.42 | 0.08 |
| **Seeds** | -0.16 | -0.23 | -0.01 | **-0.50** |
| **Other plant parts** | -0.20 | 0.02 | -0.38 | **-0.65** |
| foraging stratum: | | | | |

| | | | | |
|---|---|---|---|---|
| **Below water** | -0.02 | **0.75** | -0.03 | 0.08 |
| **Water surface** | -0.26 | **0.59** | -0.10 | -0.10 |
| **Terrestrial** | **-0.67** | -0.33 | 0.35 | -0.36 |
| **Understory** | 0.10 | -0.41 | -0.24 | 0.35 |
| **Mid to high** | **0.71** | -0.26 | -0.17 | 0.20 |
| **Canopy** | **0.73** | -0.09 | -0.04 | -0.06 |
| **Aerial** | -0.06 | -0.08 | -0.01 | 0.32 |
| **Food handling steps** | 0.04 | 0.36 | **0.78** | 0.41 |
| *SS loadings* | *2.39* | *2.32* | *2.09* | *2.00* |
| *Proportion Var* | *0.13* | *0.13* | *0.12* | *0.11* |
| *Cumulative Var* | *0.13* | *0.26* | *0.38* | *0.49* |
| *Proportion Explained* | *0.27* | *0.26* | *0.24* | *0.23* |
| *Cumulative Proportion* | *0.27* | *0.54* | *0.77* | *1.00* |

| C) Sociality | **sPC1 (Bonding)** | **sPC2 (Grouping)** |
|---|---|---|
| **Number of caretakers (N = 0-12.5)** | **0.86** | 0.00 |
| **Social bonds (short vs breeding season vs long)** | **0.86** | 0.05 |
| **Colonial breeding (yes vs no)** | -0.04 | **0.79** |
| **Grouping (non-social vs pair vs small vs large groups)** | 0.05 | **0.79** |
| *SS loadings* | *1.48* | *1.25* |
| *Proportion Var* | *0.37* | *0.31* |
| *Cumulative Var* | *0.37* | *0.68* |
| *Proportion Explained* | *0.54* | *0.46* |
| *Cumulative Proportion* | *0.54* | *1.00* |

**Table S2.** Phylogenetically controlled mixed model in the R package MCMCglmm[2] assessing the effect of relative skull volume on the relationship between mPC1-3 and brain size.

| Brain mm3 (Residual), N = 667 | Posterior Mean | lower ▪ upper 95% CI | N | p MCMC |
|---|---|---|---|---|
| Intercept | -0.384 | -0.686 ▪ -0.057 | 890 | 0.012 |
| Body mass (log-10) | 0.025 | -0.029 ▪ 0.079 | 1000 | 0.340 |
| mPC1 (Locomotion) | 0.023 | -0.030 ▪ 0.071 | 918 | 0.404 |
| **mPC2 (Beak thickness)** | **0.220** | **0.158 ▪ 0.288** | **841** | **<0.001** |
| **mPC3 (Long beak & leg)** | **0.078** | **0.026 ▪ 0.135** | **1000** | **0.006** |
| **Skull volume (residual)** | **0.312** | **0.267 ▪ 0.359** | **1000** | **<0.001** |

**Table S3**. Phylogenetically controlled mixed models in the R package MCMCglmm[2] assessing the effect on relative brain size in birds (N=1,155 species) of a) morphological predictors, b) ecological and social predictors, c) life-history predictors, and d) all combined.

| | Predictor | Posterior Mean | lower ▪ upper 95% CI | N | p MCMC |
|---|---|---|---|---|---|
| **a) Morphology model** | Intercept | -0.448 | -0.834 ▪ -0.021 | 750 | 0.022 |
| | Body mass (log-10) | 0.046 | -0.018 ▪ 0.114 | 907 | 0.188 |
| | **mPC1 (Locomotion)** | **0.077** | **0.023 ▪ 0.133** | **1107** | **0.006** |
| | **mPC2 (Beak thickness)** | **0.291** | **0.240 ▪ 0.347** | **1000** | **<0.001** |
| | **mPC3 (Long beak & leg)** | **0.139** | **0.083 ▪ 0.193** | **1000** | **<0.001** |
| **b) Eco-social model** | Intercept | -0.616 | -1.090 ▪ -0.170 | 1101 | 0.010 |
| | **Body mass (log-10)** | **0.114** | **0.049 ▪ 0.182** | **1000** | **<0.001** |
| | ePC1 (Arboreal frugivory) | 0.045 | 0.006 ▪ 0.084 | 689 | 0.020 |
| | ePC2 (Aquatic predation) | 0.014 | -0.043 ▪ 0.079 | 1000 | 0.674 |
| | ePC3 (Land predation) | 0.038 | -0.011 ▪ 0.088 | 1000 | 0.140 |
| | ePC4 (Insectivory) | 0.015 | -0.03 ▪ 0.060 | 1000 | 0.510 |
| | **sPC1 (Bonding)** | **0.048** | **0.012 ▪ 0.079** | **1000** | **0.006** |
| | sPC2 (Grouping) | -0.016 | -0.048 ▪ 0.020 | 1000 | 0.374 |
| **c) Life-history model** | Intercept | -0.631 | -1.118 ▪ -0.165 | 1000 | 0.012 |
| | Body mass (log-10) | 0.058 | -0.009 ▪ 0.125 | 894 | 0.108 |
| | **Parental provisioning (log +1)** | **0.269** | **0.168 ▪ 0.377** | **1000** | **<0.001** |
| | **Generation length (residual)** | **0.087** | **0.041 ▪ 0.128** | **1022** | **<0.001** |
| | **Migration (migratory vs sedentary)** | **0.100** | **0.040 ▪ 0.155** | **1000** | **<0.001** |
| **d) Combined model** | Intercept | -0.454 | -0.836 ▪ -0.033 | 1000 | 0.030 |
| | Body mass (log-10) | 0.014 | -0.058 ▪ 0.075 | 1000 | 0.680 |
| | mPC1 (Locomotion) | 0.064 | 0.009 ▪ 0.115 | 1000 | 0.014 |
| | **mPC2 (Beak thickness)** | **0.272** | **0.215 ▪ 0.329** | **1000** | **<0.001** |
| | **mPC3 (Long beak & leg)** | **0.125** | **0.076 ▪ 0.185** | **1000** | **<0.001** |
| | **Parental providing (log +1)** | **0.171** | **0.080 ▪ 0.268** | **992** | **<0.001** |
| | **Generation length (residual)** | **0.086** | **0.042 ▪ 0.127** | **1000** | **<0.001** |
| | **Migration (migratory vs sedentary)** | **0.087** | **0.031 ▪ 0.147** | **1197** | **0.002** |
| | ePC1 (Arboreal frugivory) | 0.005 | -0.031 ▪ 0.045 | 1000 | 0.804 |
| | ePC2 (Aquatic predation) | -0.004 | -0.066 ▪ 0.055 | 1000 | 0.868 |
| | ePC3 (Land Predation) | 0.036 | -0.010 ▪ 0.080 | 1000 | 0.126 |
| | ePC4 (Insectivory) | 0.004 | -0.035 ▪ 0.045 | 1000 | 0.898 |
| | sPC1 (Bonding) | 0.015 | -0.016 ▪ 0.048 | 1000 | 0.334 |
| | sPC2 (Grouping) | -0.018 | -0.052 ▪ 0.009 | 1000 | 0.264 |

**Table S4.** Phylogenetically controlled mixed models in the r package MCMCglmm[2] assessing the effect on relative brain size in birds (N= 660 species with data on eye size) of a) morphology predictors, b) eye size, c) eco-social predictors, d) life-history predictors, and e) all combined. The combined model explained more variance in relative brain size (R2 = 0.574) than the morphological (R2 = 0.488), life-history (R2 = 0.438), relative eye size (R2 = 0.242) or eco-social model (R2 = 0.347).

| | Predictor | Posterior Mean | lower ▪ upper 95% CI | N | p MCMC |
|---|---|---|---|---|---|
| **a) Morphology model** | Intercept | -0.482 | -0.881 ▪ -0.089 | 1000 | 0.018 |
| | Body mass (log-10) | 0.092 | 0.019 ▪ 0.171 | 997 | 0.030 |
| | **mPC1 (Locomotion)** | **0.089** | **0.038 ▪ 0.151** | **1000** | **0.006** |
| | **mPC2 (Beak thickness)** | **0.304** | **0.240 ▪ 0.375** | **1000** | **<0.001** |
| | **mPC3 (Long beak & leg)** | **0.126** | **0.067 ▪ 0.183** | **1000** | **<0.001** |
| **b) Eye size** | Intercept | -0.728 | -1.144 ▪ -0.242 | 1069 | <0.001 |
| | **Body mass (log-10)** | **0.171** | **0.090 ▪ 0.251** | **1000** | **<0.001** |
| | **Eye size (residual)** | **0.121** | **0.071 ▪ 0.173** | **1000** | **<0.001** |
| **c) Eco-social model** | Intercept | -0.721 | -1.179 ▪ -0.287 | 903 | 0.001 |
| | **Body mass (log-10)** | 0.195 | 0.109 ▪ 0.268 | 1000 | <0.001 |
| | **ePC1 (Arboreal frugivory)** | **0.062** | **0.015 ▪ 0.103** | **900** | **0.006** |
| | ePC2 (Aquatic predation) | -0.039 | -0.095 ▪ 0.025 | 1229 | 0.200 |
| | ePC3 (Land Predation) | 0.000 | -0.063 ▪ 0.057 | 1229 | 0.970 |
| | ePC4 (Insectivory) | 0.008 | -0.041 ▪ 0.068 | 1000 | 0.792 |
| | **sPC1 (Bonding)** | **0.080** | **0.035 ▪ 0.124** | **1000** | **0.002** |
| | sPC2 (Grouping) | -0.036 | -0.076 ▪ 0.006 | 1000 | 0.080 |
| **d) Life-history model** | Intercept | -0.676 | -1.156 ▪ -0.254 | 1000 | 0.002 |
| | **Body mass (log-10)** | **0.119** | **0.041 ▪ 0.192** | **1000** | **0.004** |
| | **Parental providing (log +1)** | **0.244** | **0.130 ▪ 0.363** | **1000** | **<0.001** |
| | **Generation length (residual)** | **0.077** | **0.033 ▪ 0.124** | **1000** | **<0.001** |
| | **Migration (migratory vs sedentary)** | **0.112** | **0.037 ▪ 0.180** | **1000** | **0.004** |
| **e) Combined model** | Intercept | -0.463 | -0.811 ▪ -0.072 | 1000 | 0.010 |
| | Body mass (log-10) | 0.087 | 0.014 ▪ 0.174 | 907 | 0.038 |
| | **Eye size (residual)** | **0.069** | **0.017 ▪ 0.120** | **1000** | **0.004** |
| | mPC1 (Locomotion) | 0.062 | 0.008 ▪ 0.125 | 1000 | 0.042 |
| | **mPC2 (Beak thickness)** | **0.261** | **0.195 ▪ 0.335** | **1543** | **<0.001** |
| | **mPC3 (Long beak & leg)** | **0.105** | **0.039 ▪ 0.165** | **1000** | **0.002** |
| | **Parental providing (log +1)** | **0.155** | **0.046 ▪ 0.263** | **1000** | **0.008** |
| | **Generation length (residual)** | **0.062** | **0.021 ▪ 0.106** | **1138** | **0.004** |
| | Migration (migratory vs sedentary) | 0.094 | 0.022 ▪ 0.163 | 1000 | 0.014 |
| | ePC1 (Arboreal frugivory) | 0.007 | -0.034 ▪ 0.052 | 1093 | 0.750 |
| | ePC2 (Aquatic predation) | -0.026 | -0.080 ▪ 0.026 | 1000 | 0.360 |
| | ePC3 (Land Predation) | -0.006 | -0.061 ▪ 0.050 | 1000 | 0.860 |
| | ePC4 (Insectivory) | 0.002 | -0.049 ▪ 0.057 | 1150 | 0.928 |
| | sPC1 (Bonding) | 0.035 | -0.009 ▪ 0.076 | 1000 | 0.112 |
| | sPC2 (Grouping) | -0.029 | -0.065 ▪ 0.008 | 1000 | 0.134 |

**Table S5.** Phylogenetically controlled mixed models in the r package MCMCglmm[2] assessing the effect on brain region including mass and neuron numbers of telencephalon, pallium, cerebellum and brainstem (n =110) of morphology predictors (mPC1-3).

| | Posterior Mean | lower ▪ upper 95% CI | N | p MCMC |
|---|---|---|---|---|
| **mPC1 (Locomotion) effects on brain regions** | | | | |
| **Telencephalon mass (residual)** | | | | |
| (Intercept) | -0.994 | -1.661 ▪ -0.294 | 1000 | 0.004 |
| Body mass (log-10) | 0.271 | 0.155 ▪ 0.374 | 1286 | <0.001 |
| Development mode (*altricial*) | 1.335 | 0.448 ▪ 2.215 | 1000 | 0.006 |
| mPC1 | 0.061 | -0.044 ▪ 0.171 | 1000 | 0.256 |
| **Telencephalon neuron numbers (residual)** | | | | |
| (Intercept) | -0.450 | -0.731 ▪ -0.198 | 1000 | 0.002 |
| Body mass (log-10) | 0.142 | 0.095 ▪ 0.191 | 1000 | <0.001 |
| Development mode (*altricial*) | 0.619 | 0.259 ▪ 0.990 | 1000 | <0.001 |
| mPC1 | 0.005 | -0.044 ▪ 0.058 | 1000 | 0.852 |
| **Pallium mass (residual)** | | | | |
| (Intercept) | -1.001 | -1.662 ▪ -0.361 | 1000 | 0.004 |
| Body mass (log-10) | 0.260 | 0.161 ▪ 0.366 | 1000 | <0.001 |
| Development mode (*altricial*) | 1.336 | 0.419 ▪ 2.228 | 1000 | 0.002 |
| mPC1 | 0.047 | -0.054 ▪ 0.148 | 1000 | 0.396 |
| **Pallium neuron numbers (residual)** | | | | |
| (Intercept) | -0.458 | -0.741 ▪ -0.198 | 1000 | 0.002 |
| Body mass (log-10) | 0.136 | 0.087 ▪ 0.188 | 1000 | <0.001 |
| Development mode (*altricial*) | 0.625 | 0.214 ▪ 0.969 | 1000 | 0.006 |
| mPC1 | 0.004 | -0.044 ▪ 0.065 | 1058 | 0.902 |
| **Cerebellum mass (residual)** | | | | |
| (Intercept) | -0.946 | -1.630 ▪ -0.114 | 1000 | 0.016 |
| Body mass (log-10) | 0.194 | 0.029 ▪ 0.349 | 1000 | 0.026 |
| Development mode (*altricial*) | 1.391 | 0.401 ▪ 2.499 | 1000 | 0.014 |
| **mPC1** | **0.149** | **-0.005 ▪ 0.296** | **1210** | **0.044** |
| **Cerebellum neuron numbers (residual)** | | | | |

| | | | | |
|---|---|---|---|---|
| (Intercept) | -0.194 | -0.331 ▪ -0.060 | 1000 | 0.010 |
| Body mass (log-10) | 0.069 | 0.039 ▪ 0.100 | 1000 | <0.001 |
| Development mode (*altricial*) | 0.287 | 0.102 ▪ 0.467 | 1000 | <0.001 |
| **mPC1** | **0.033** | **0.003 ▪ 0.064** | **1000** | **0.040** |
| **Brainstem mass (residual)** | | | | |
| (Intercept) | -0.969 | -1.845 ▪ -0.026 | 874 | 0.044 |
| Body mass (log-10) | 0.173 | 0.025 ▪ 0.356 | 1000 | 0.038 |
| Development mode (*altricial*) | 1.359 | 0.189 ▪ 2.660 | 1000 | 0.030 |
| mPC1 | 0.082 | -0.097 ▪ 0.261 | 1000 | 0.364 |
| **Brainstem neuron numbers (residual)** | | | | |
| (Intercept) | -0.121 | -0.241 ▪ 0.003 | 1000 | 0.044 |
| Body mass (log-10) | 0.033 | -0.005 ▪ 0.076 | 1000 | 0.100 |
| Development mode (*altricial*) | 0.201 | 0.034 ▪ 0.362 | 1000 | 0.016 |
| mPC1 | -0.001 | -0.045 ▪ 0.048 | 1000 | 0.970 |
| **mPC2 (Beak thickness) effects on brain regions** | | | | |
| **Telencephalon mass (residual)** | | | | |
| (Intercept) | -0.939 | -1.505 ▪ -0.289 | 1293 | 0.002 |
| Body mass (log-10) | 0.311 | 0.225 ▪ 0.407 | 1000 | <0.001 |
| Development mode (*altricial*) | 1.335 | 0.593 ▪ 2.213 | 1000 | <0.001 |
| **mPC2** | **0.191** | **0.036 ▪ 0.321** | **1000** | **0.010** |
| **Telencephalon neuron numbers (residual)** | | | | |
| (Intercept) | -0.390 | -0.661 ▪ -0.122 | 1000 | <0.001 |
| Body mass (log-10) | 0.146 | 0.103 ▪ 0.183 | 1000 | <0.001 |
| Development mode (*altricial*) | 0.563 | 0.228 ▪ 0.916 | 1000 | <0.001 |
| **mPC2** | **0.093** | **0.032 ▪ 0.156** | **1000** | **0.008** |
| **Pallium mass (residual)** | | | | |
| (Intercept) | -0.935 | -1.460 ▪ -0.330 | 856 | 0.004 |
| Body mass (log-10) | 0.291 | 0.200 ▪ 0.381 | 1000 | <0.001 |
| Development mode (*altricial*) | 1.317 | 0.514 ▪ 2.088 | 1000 | <0.001 |
| **mPC2** | **0.176** | **0.052 ▪ 0.341** | **1000** | **0.018** |
| **Pallium neuron numbers (residual)** | | | | |

| | | | | |
|---|---|---|---|---|
| (Intercept) | -0.401 | -0.667 ▪ -0.159 | 1000 | 0.002 |
| Body mass (log-10) | 0.139 | 0.099 ▪ 0.180 | 1004 | <0.001 |
| Development mode (*altricial*) | 0.575 | 0.247 ▪ 0.890 | 1316 | <0.001 |
| **mPC2** | **0.087** | **0.022 ▪ 0.154** | **1000** | **0.010** |
| **Cerebellum mass (residual)** | | | | |
| (Intercept) | -1.051 | -1.709 ▪ -0.411 | 1000 | 0.002 |
| Body mass (log-10) | 0.272 | 0.148 ▪ 0.407 | 1000 | <0.001 |
| Development mode (*altricial*) | 1.595 | 0.748 ▪ 2.509 | 1000 | 0.002 |
| mPC2 | 0.133 | -0.097 ▪ 0.319 | 1000 | 0.224 |
| **Cerebellum neuron numbers (residual)** | | | | |
| (Intercept) | -0.214 | -0.334 ▪ -0.084 | 977 | 0.004 |
| Body mass (log-10) | 0.084 | 0.058 ▪ 0.111 | 1000 | <0.001 |
| Development mode (*altricial*) | 0.330 | 0.157 ▪ 0.495 | 1107 | <0.001 |
| mPC2 | 0.033 | -0.006 ▪ 0.075 | 1103 | 0.118 |
| **Brainstem mass (residual)** | | | | |
| (Intercept) | -0.980 | -1.806 ▪ -0.207 | 1000 | 0.028 |
| Body mass (log-10) | 0.223 | 0.074 ▪ 0.375 | 1000 | 0.004 |
| Development mode (*altricial*) | 1.419 | 0.425 ▪ 2.540 | 993 | 0.006 |
| mPC2 | 0.173 | -0.051 ▪ 0.412 | 1000 | 0.144 |
| **Brainstem neuron numbers (residual)** | | | | |
| (Intercept) | -0.111 | -0.217 ▪ 0.012 | 1000 | 0.066 |
| Body mass (log-10) | 0.035 | -0.002 ▪ 0.066 | 1000 | 0.052 |
| Development mode (*altricial*) | 0.192 | 0.051 ▪ 0.331 | 1093 | 0.010 |
| mPC2 | 0.019 | -0.023 ▪ 0.067 | 1000 | 0.396 |
| **mPC3 (Long beak & leg) effects on brain regions** | | | | |
| **Telencephalon mass (residual)** | | | | |
| (Intercept) | -1.092 | -1.796 ▪ -0.446 | 1000 | 0.002 |
| Body mass (log-10) | 0.309 | 0.212 ▪ 0.420 | 1000 | <0.001 |
| Development mode (*altricial*) | 1.491 | 0.515 ▪ 2.339 | 1000 | 0.004 |
| mPC3 | -0.027 | -0.142 ▪ 0.078 | 1105 | 0.612 |
| **Telencephalon neuron numbers (residual)** | | | | |

| | | | | |
|---|---|---|---|---|
| (Intercept) | -0.457 | -0.729 ▪ -0.172 | 1000 | 0.004 |
| Body mass (log-10) | 0.136 | 0.088 ▪ 0.186 | 1000 | <0.001 |
| Development mode (*altricial*) | 0.618 | 0.284 ▪ 0.988 | 858 | 0.004 |
| mPC3 | 0.021 | -0.026 ▪ 0.073 | 903 | 0.404 |
| **Pallium mass (residual)** | | | | |
| (Intercept) | -1.052 | -1.632 ▪ -0.375 | 1000 | <0.001 |
| Body mass (log-10) | 0.300 | 0.195 ▪ 0.397 | 1000 | <0.001 |
| Development mode (*altricial*) | 1.452 | 0.627 ▪ 2.343 | 1000 | <0.001 |
| mPC3 | -0.047 | -0.153 ▪ 0.056 | 898 | 0.386 |
| **Pallium neuron numbers (residual)** | | | | |
| (Intercept) | -0.456 | -0.706 ▪ -0.149 | 818 | 0.002 |
| Body mass (log-10) | 0.130 | 0.080 ▪ 0.179 | 1000 | <0.001 |
| Development mode (*altricial*) | 0.615 | 0.229 ▪ 0.951 | 1000 | <0.001 |
| mPC3 | 0.021 | -0.029 ▪ 0.071 | 1118 | 0.404 |
| **Cerebellum mass (residual)** | | | | |
| (Intercept) | -1.123 | -1.810 ▪ -0.448 | 1000 | 0.006 |
| Body mass (log-10) | 0.252 | 0.086 ▪ 0.379 | 1000 | <0.001 |
| Development mode (*altricial*) | 1.661 | 0.730 ▪ 2.568 | 1000 | 0.008 |
| mPC3 | 0.016 | -0.140 ▪ 0.147 | 1000 | 0.856 |
| **Cerebellum neuron numbers (residual)** | | | | |
| (Intercept) | -0.237 | -0.386 ▪ -0.098 | 1000 | <0.001 |
| Body mass (log-10) | 0.073 | 0.046 ▪ 0.103 | 880 | <0.001 |
| Development mode (*altricial*) | 0.335 | 0.153 ▪ 0.511 | 1000 | <0.001 |
| mPC3 | 0.024 | -0.008 ▪ 0.053 | 1000 | 0.110 |
| **Brainstem mass (residual)** | | | | |
| (Intercept) | -1.121 | -2.093 ▪ -0.358 | 1102 | 0.022 |
| Body mass (log-10) | 0.196 | 0.049 ▪ 0.372 | 1000 | 0.024 |
| Development mode (*altricial*) | 1.538 | 0.454 ▪ 2.801 | 1000 | 0.010 |
| mPC3 | 0.036 | -0.134 ▪ 0.181 | 1000 | 0.638 |
| **Brainstem neuron numbers (residual)** | | | | |
| (Intercept) | -0.117 | -0.219 ▪ -0.009 | 1000 | 0.030 |

| | | | | |
|---|---|---|---|---|
| Body mass (log-10) | 0.031 | -0.006 ▪ 0.068 | 1355 | 0.114 |
| Development mode (*altricial*) | 0.195 | 0.051 ▪ 0.331 | 1000 | 0.016 |
| mPC3 | 0.006 | -0.031 ▪ 0.045 | 1115 | 0.786 |

**Table S6**. Phylogenetic Generalized Least Squares (PGLS) models in the package phylolm[3] showing the association between brain size, morphology (mPC1-3), eco-social niche (ePC1-4, sPC1-2) and life-history traits based on 1155 species.

| Predictor | Estimate ± se | t value | p value |
|---|---|---|---|
| **Model: brain ~ other traits** | | | |
| (Intercept) | -0.405 ± 0.203 | -1.997 | 0.046 |
| Body mass (log-10) | 0.014 ± 0.034 | 0.412 | 0.680 |
| **mPC1 (Locomotion)** | 0.063 ± 0.028 | 2.284 | 0.023 |
| **mPC2 (Beak thickness)** | 0.274 ± 0.029 | 9.592 | <0.001 |
| **mPC3 (Long beak & leg)** | 0.125 ± 0.028 | 4.493 | <0.001 |
| ePC1 (Arboreal frugivory) | 0.006 ± 0.019 | 0.309 | 0.757 |
| ePC2 (Aquatic predation) | -0.006 ± 0.030 | -0.183 | 0.855 |
| ePC3 (Land Predation) | 0.035 ± 0.024 | 1.484 | 0.138 |
| ePC4 (Insectivory) | 0.004 ± 0.021 | 0.190 | 0.850 |
| sPC1 (Bonding) | 0.016 ± 0.016 | 0.959 | 0.338 |
| sPC2 (Grouping) | -0.017 ± 0.017 | -1.040 | 0.298 |
| **Parental providing (log +1)** | 0.171 ± 0.051 | 3.332 | <0.001 |
| **Generation length (residual)** | 0.085 ± 0.022 | 3.955 | <0.001 |
| **Migration (migratory vs sedentary)** | -0.044 ± 0.015 | -2.963 | 0.003 |
| | | lambda = 0.867; r2 = 0.179 | |
| **Model: mPC1 ~ other traits** | | | |
| (Intercept) | -0.270 ± 0.285 | -0.945 | 0.345 |
| Body mass (log-10) | 0.242 ± 0.038 | 6.446 | <0.001 |
| Brain size (residual) | 0.047 ± 0.029 | 1.607 | 0.108 |
| ePC1 (Arboreal frugivory) | 0.021 ± 0.020 | 1.025 | 0.305 |
| **ePC2 (Aquatic predation)** | -0.198 ± 0.031 | -6.308 | <0.001 |
| ePC3 (Land Predation) | 0.040 ± 0.025 | 1.585 | 0.113 |
| ePC4 (Insectivory) | -0.043 ± 0.022 | -1.955 | 0.051 |
| sPC1 (Bonding) | 0.001 ± 0.017 | 0.034 | 0.973 |
| sPC2 (Grouping) | -0.009 ± 0.017 | -0.560 | 0.575 |
| Parental providing (log +1) | -0.101 ± 0.056 | -1.820 | 0.069 |
| **Generation length (residual)** | 0.065 ± 0.022 | 2.913 | 0.004 |
| Migration (migratory vs sedentary) | -0.003 ± 0.015 | -0.191 | 0.849 |
| | | lambda = 0.963; r2 = 0.086 | |
| **Model: mPC2 ~ other traits** | | | |
| (Intercept) | -0.281 ± 0.255 | -1.100 | 0.272 |
| Body mass (log-10) | 0.065 ± 0.035 | 1.846 | 0.065 |
| **Brain size (residual)** | 0.265 ± 0.028 | 9.457 | <0.001 |

| | | | |
|---|---|---|---|
| **ePC1 (Arboreal frugivory)** | 0.089 ± 0.019 | 4.569 | <0.001 |
| ePC2 (Aquatic predation) | 0.015 ± 0.030 | 0.494 | 0.621 |
| ePC3 (Land Predation) | -0.042 ± 0.024 | -1.755 | 0.080 |
| ePC4 (Insectivory) | 0.024 ± 0.021 | 1.138 | 0.255 |
| **sPC1 (Bonding)** | 0.036 ± 0.017 | 2.140 | 0.033 |
| sPC2 (Grouping) | 0.025 ± 0.017 | 1.521 | 0.129 |
| **Parental providing (log +1)** | 0.234 ± 0.053 | 4.385 | <0.001 |
| Generation length (residual) | -0.038 ± 0.022 | -1.754 | 0.080 |
| **Migration (migratory vs sedentary)** | 0.033 ± 0.014 | 2.301 | 0.022 |
| | | lambda = 0.946; r2 = 0.139 | |
| **Model: mPC3 ~ other traits** | | | |
| (Intercept) | -0.131 ± 0.273 | -0.478 | 0.633 |
| **Body mass (log-10)** | 0.210 ± 0.037 | 5.748 | <0.001 |
| **Brain size (residual)** | 0.208 ± 0.028 | 7.302 | <0.001 |
| ePC1 (Arboreal frugivory) | -0.017 ± 0.020 | -0.847 | 0.397 |
| ePC2 (Aquatic predation) | 0.050 ± 0.031 | 1.622 | 0.105 |
| ePC3 (Land Predation) | 0.041 ± 0.025 | 1.683 | 0.093 |
| ePC4 (Insectivory) | 0.029 ± 0.021 | 1.363 | 0.173 |
| sPC1 (Bonding) | -0.016 ± 0.017 | -0.924 | 0.356 |
| sPC2 (Grouping) | -0.003 ± 0.017 | -0.167 | 0.868 |
| Parental providing (log +1) | -0.021 ± 0.055 | -0.378 | 0.705 |
| Generation length (residual) | -0.015 ± 0.022 | -0.661 | 0.508 |
| **Migration (migratory vs sedentary)** | -0.050 ± 0.015 | -3.424 | <0.001 |
| | | lambda = 0.958; r2 = 0.100 | |
| **Model: ePC1 ~ other traits** | | | |
| (Intercept) | 0.025 ± 0.447 | 0.057 | 0.955 |
| Body mass (log-10) | 0.025 ± 0.058 | 0.423 | 0.673 |
| Brain size (residual) | 0.027 ± 0.046 | 0.583 | 0.560 |
| mPC1 (Locomotion) | -0.015 ± 0.046 | -0.319 | 0.750 |
| **mPC2 (Beak thickness)** | 0.241 ± 0.049 | 4.908 | <0.001 |
| mPC3 (Long beak & leg) | -0.052 ± 0.048 | -1.083 | 0.279 |
| **sPC1 (Bonding)** | -0.081 ± 0.027 | -3.049 | 0.002 |
| sPC2 (Grouping) | -0.024 ± 0.026 | -0.929 | 0.353 |
| **Parental providing (log +1)** | 0.170 ± 0.086 | 1.983 | 0.048 |
| **Generation length (residual)** | 0.081 ± 0.034 | 2.383 | 0.017 |
| Migration (migratory vs sedentary) | -0.021 ± 0.023 | -0.928 | 0.354 |
| | | lambda = 0.968; r2 = 0.046 | |
| **Model: ePC2 ~ other traits** | | | |

| | | | |
|---|---|---|---|
| (Intercept) | -0.051 ± 0.235 | -0.215 | 0.830 |
| **Body mass (log-10)** | 0.095 ± 0.035 | 2.686 | 0.007 |
| Brain size (residual) | 0.002 ± 0.030 | 0.076 | 0.939 |
| **mPC1 (Locomotion)** | -0.218 ± 0.028 | -7.699 | <0.001 |
| mPC2 (Beak thickness) | 0.020 ± 0.031 | 0.666 | 0.506 |
| **mPC3 (Long beak & leg)** | 0.071 ± 0.030 | 2.381 | 0.017 |
| **sPC1 (Bonding)** | -0.035 ± 0.017 | -2.092 | 0.037 |
| sPC2 (Grouping) | 0.009 ± 0.017 | 0.525 | 0.599 |
| Parental providing (log +1) | 0.087 ± 0.054 | 1.607 | 0.108 |
| **Generation length (residual)** | 0.101 ± 0.022 | 4.557 | <0.001 |
| Migration (migratory vs sedentary) | 0.006 ± 0.015 | 0.399 | 0.690 |
| | | lambda = 0.913; r2 = 0.083 | |
| **Model: ePC3 ~ other traits** | | | |
| (Intercept) | -0.171 ± 0.284 | -0.602 | 0.548 |
| **Body mass (log-10)** | 0.155 ± 0.043 | 3.628 | <0.001 |
| Brain size (residual) | 0.049 ± 0.037 | 1.335 | 0.182 |
| mPC1 (Locomotion) | 0.059 ± 0.034 | 1.744 | 0.081 |
| mPC2 (Beak thickness) | -0.051 ± 0.037 | -1.388 | 0.165 |
| mPC3 (Long beak & leg) | 0.059 ± 0.036 | 1.639 | 0.101 |
| **sPC1 (Bonding)** | -0.040 ± 0.020 | -1.965 | 0.050 |
| **sPC2 (Grouping)** | -0.083 ± 0.020 | -4.068 | <0.001 |
| Parental providing (log +1) | 0.053 ± 0.065 | 0.813 | 0.416 |
| Generation length (residual) | 0.000 ± 0.027 | -0.018 | 0.986 |
| Migration (migratory vs sedentary) | 0.005 ± 0.018 | 0.286 | 0.775 |
| | | lambda = 0.914; r2 = 0.047 | |
| **Model: ePC4 ~ other traits** | | | |
| (Intercept) | -0.034 ± 0.272 | -0.126 | 0.900 |
| **Body mass (log-10)** | -0.304 ± 0.046 | -6.597 | <0.001 |
| Brain size (residual) | 0.010 ± 0.042 | 0.230 | 0.818 |
| **mPC1 (Locomotion)** | -0.095 ± 0.037 | -2.567 | 0.010 |
| mPC2 (Beak thickness) | 0.025 ± 0.041 | 0.621 | 0.535 |
| **mPC3 (Long beak & leg)** | 0.080 ± 0.039 | 2.075 | 0.038 |
| sPC1 (Bonding) | -0.036 ± 0.023 | -1.585 | 0.113 |
| **sPC2 (Grouping)** | -0.075 ± 0.023 | -3.224 | 0.001 |
| Parental providing (log +1) | -0.010 ± 0.072 | -0.133 | 0.894 |
| **Generation length (residual)** | 0.080 ± 0.030 | 2.660 | 0.008 |
| Migration (migratory vs sedentary) | 0.009 ± 0.021 | 0.435 | 0.663 |
| | | lambda = 0.845; r2 = 0.067 | |

| | | | |
|---|---|---|---|
| **Model: sPC1 ~ other traits** | | | |
| (Intercept) | -0.151 ± 0.404 | -0.374 | 0.708 |
| **Body mass (log-10)** | -0.168 ± 0.063 | -2.671 | 0.008 |
| Brain size (residual) | 0.051 ± 0.053 | 0.971 | 0.332 |
| mPC1 (Locomotion) | 0.045 ± 0.051 | 0.888 | 0.374 |
| **mPC2 (Beak thickness)** | 0.132 ± 0.054 | 2.456 | 0.014 |
| mPC3 (Long beak & leg) | -0.071 ± 0.052 | -1.376 | 0.169 |
| **ePC1 (Arboreal frugivory)** | -0.077 ± 0.034 | -2.225 | 0.026 |
| ePC2 (Aquatic predation) | -0.065 ± 0.055 | -1.185 | 0.236 |
| ePC3 (Land Predation) | -0.074 ± 0.043 | -1.749 | 0.081 |
| ePC4 (Insectivory) | -0.036 ± 0.038 | -0.950 | 0.343 |
| **Parental providing (log +1)** | 0.281 ± 0.094 | 2.992 | 0.003 |
| Generation length (residual) | 0.039 ± 0.039 | 1.015 | 0.310 |
| **Migration (migratory vs sedentary)** | -0.126 ± 0.026 | -4.847 | <0.001 |
| | | lambda = 0.910; r2 = 0.056 | |
| **Model: sPC2 ~ other traits** | | | |
| (Intercept) | -0.237 ± 0.287 | -0.828 | 0.408 |
| Body mass (log-10) | 0.100 ± 0.056 | 1.782 | 0.075 |
| Brain size (residual) | -0.051 ± 0.051 | -0.989 | 0.323 |
| mPC1 (Locomotion) | 0.014 ± 0.046 | 0.307 | 0.759 |
| mPC2 (Beak thickness) | 0.088 ± 0.049 | 1.779 | 0.076 |
| mPC3 (Long beak & leg) | -0.052 ± 0.046 | -1.130 | 0.259 |
| ePC1 (Arboreal frugivory) | -0.032 ± 0.032 | -1.003 | 0.316 |
| ePC2 (Aquatic predation) | 0.088 ± 0.051 | 1.714 | 0.087 |
| **ePC3 (Land Predation)** | -0.171 ± 0.040 | -4.249 | <0.001 |
| **ePC4 (Insectivory)** | -0.115 ± 0.037 | -3.103 | 0.002 |
| Parental providing (log +1) | 0.014 ± 0.085 | 0.167 | 0.867 |
| Generation length (residual) | 0.054 ± 0.038 | 1.431 | 0.153 |
| **Migration (migratory vs sedentary)** | 0.062 ± 0.026 | 2.382 | 0.017 |
| | | lambda = 0.756; r2 = 0.044 | |
| **Model: Time fed ~ other traits** | | | |
| (Intercept) | -0.144 ± 0.141 | -1.018 | 0.309 |
| **Body mass (log-10)** | 0.169 ± 0.020 | 8.685 | <0.001 |
| **Brain size (residual)** | 0.049 ± 0.016 | 3.032 | 0.002 |
| mPC1 (Locomotion) | -0.011 ± 0.016 | -0.663 | 0.507 |
| **mPC2 (Beak thickness)** | 0.073 ± 0.017 | 4.308 | <0.001 |
| mPC3 (Long beak & leg) | -0.020 ± 0.016 | -1.219 | 0.223 |
| ePC1 (Arboreal frugivory) | 0.019 ± 0.011 | 1.707 | 0.088 |

| | | | |
|---|---|---|---|
| ePC2 (Aquatic predation) | 0.014 ± 0.017 | 0.841 | 0.400 |
| ePC3 (Land Predation) | 0.010 ± 0.013 | 0.734 | 0.463 |
| ePC4 (Insectivory) | -0.013 ± 0.012 | -1.113 | 0.266 |
| **sPC1 (Bonding)** | 0.025 ± 0.009 | 2.740 | 0.006 |
| sPC2 (Grouping) | 0.000 ± 0.009 | 0.031 | 0.976 |
| **Generation length (residual)** | 0.040 ± 0.012 | 3.344 | <0.001 |
| **Migration (migratory vs sedentary)** | -0.038 ± 0.008 | -4.828 | <0.001 |
| | | lambda = 0.947; r2 = 0.159 | |
| **Model: Generation length ~ other traits** | | | |
| (Intercept) | -0.024 ± 0.270 | -0.087 | 0.931 |
| Body mass (log-10) | -0.009 ± 0.046 | -0.195 | 0.846 |
| **Brain size (residual)** | 0.159 ± 0.040 | 3.976 | <0.001 |
| **mPC1 (Locomotion)** | 0.134 ± 0.037 | 3.594 | <0.001 |
| mPC2 (Beak thickness) | -0.057 ± 0.040 | -1.426 | 0.154 |
| mPC3 (Long beak & leg) | -0.031 ± 0.038 | -0.810 | 0.418 |
| ePC1 (Arboreal frugivory) | 0.010 ± 0.026 | 0.398 | 0.691 |
| **ePC2 (Aquatic predation)** | 0.159 ± 0.041 | 3.896 | <0.001 |
| ePC3 (Land Predation) | 0.000 ± 0.032 | -0.008 | 0.994 |
| **ePC4 (Insectivory)** | 0.058 ± 0.029 | 1.997 | 0.046 |
| sPC1 (Bonding) | 0.024 ± 0.022 | 1.056 | 0.291 |
| sPC2 (Grouping) | 0.030 ± 0.023 | 1.296 | 0.195 |
| **Parental providing (log +1)** | 0.275 ± 0.069 | 3.969 | <0.001 |
| Migration (migratory vs sedentary) | 0.016 ± 0.020 | 0.809 | 0.419 |
| | | lambda = 0.856; r2 = 0.067 | |

**Table S7**. Phylogenetic Generalized Least Squares (PGLS) models in the package phylolm[3] the association between brain size, eye size, morphology (mPC1-3), eco-social niche (ePC1-4 and sPC1-2) and life-history traits based on 660 species where data on eye size was available.

| | Estimate | t value | p value |
|---|---|---|---|
| **Model: brain ~ other traits** | | | |
| (Intercept) | -0.407 ± 0.196 | -2.080 | 0.038 |
| **Body mass (log-10)** | 0.088 ± 0.040 | 2.215 | 0.027 |
| **Eye size (residual)** | 0.732 ± 0.281 | 2.602 | 0.009 |
| **mPC1 (Locomotion)** | 0.061 ± 0.030 | 2.052 | 0.041 |
| **mPC2 (Beak thickness)** | 0.261 ± 0.035 | 7.422 | <0.001 |
| **mPC3 (Long beak & leg)** | 0.104 ± 0.031 | 3.312 | <0.001 |
| ePC1 (Arboreal frugivory) | 0.006 ± 0.022 | 0.286 | 0.775 |
| ePC2 (Aquatic predation) | -0.025 ± 0.028 | -0.901 | 0.368 |
| ePC3 (Land Predation) | -0.006 ± 0.028 | -0.225 | 0.822 |
| ePC4 (Insectivory) | 0.001 ± 0.027 | 0.045 | 0.964 |
| sPC1 (Bonding) | 0.034 ± 0.022 | 1.570 | 0.117 |
| sPC2 (Grouping) | -0.029 ± 0.019 | -1.516 | 0.130 |
| **Parental providing (log +1)** | 0.153 ± 0.055 | 2.768 | 0.006 |
| **Generation length (residual)** | 0.064 ± 0.022 | 2.895 | 0.004 |
| **Migration (migratory vs sedentary)** | -0.047 ± 0.018 | -2.570 | 0.010 |
| | | lambda = 0.888; r2 = 0.232 | |
| **Model: Eye ~ other traits** | | | |
| (Intercept) | -0.004 ± 0.018 | -0.252 | 0.801 |
| Body mass (log-10) | -0.003 ± 0.005 | -0.541 | 0.588 |
| **Brain size (residual)** | 0.013 ± 0.005 | 2.640 | 0.008 |
| **mPC1 (Locomotion)** | 0.019 ± 0.004 | 5.142 | <0.001 |
| **mPC2 (Beak thickness)** | 0.012 ± 0.005 | 2.643 | 0.008 |
| mPC3 (Long beak & leg) | -0.005 ± 0.004 | -1.211 | 0.226 |
| ePC1 (Arboreal frugivory) | 0.003 ± 0.003 | 1.146 | 0.252 |
| ePC2 (Aquatic predation) | -0.003 ± 0.003 | -0.828 | 0.408 |
| **ePC3 (Land Predation)** | 0.018 ± 0.004 | 5.003 | <0.001 |
| **ePC4 (Insectivory)** | 0.015 ± 0.003 | 4.318 | <0.001 |
| sPC1 (Bonding) | 0.002 ± 0.003 | 0.853 | 0.394 |
| sPC2 (Grouping) | -0.005 ± 0.003 | -1.921 | 0.055 |
| Parental providing (log +1) | 0.003 ± 0.007 | 0.378 | 0.705 |
| Generation length (residual) | 0.000 ± 0.003 | 0.065 | 0.948 |
| Migration (migratory vs sedentary) | -0.002 ± 0.003 | -0.722 | 0.471 |
| | | lambda = 0.638; r2 = 0.164 | |

| | | | |
|---|---|---|---|
| **Model: mPC1 ~ other traits** | | | |
| (Intercept) | -0.574 ± 0.313 | -1.830 | 0.068 |
| **Body mass (log-10)** | 0.352 ± 0.052 | 6.775 | <0.001 |
| Brain size (residual) | 0.049 ± 0.049 | 0.997 | 0.319 |
| **Eye size (residual)** | 1.609 ± 0.346 | 4.645 | <0.001 |
| ePC1 (Arboreal frugivory) | -0.014 ± 0.028 | -0.491 | 0.624 |
| **ePC2 (Aquatic predation)** | -0.107 ± 0.038 | -2.819 | 0.005 |
| ePC3 (Land Predation) | 0.070 ± 0.037 | 1.888 | 0.059 |
| ePC4 (Insectivory) | 0.039 ± 0.035 | 1.119 | 0.263 |
| sPC1 (Bonding) | -0.038 ± 0.029 | -1.328 | 0.185 |
| sPC2 (Grouping) | 0.001 ± 0.024 | 0.060 | 0.952 |
| Parental providing (log +1) | -0.031 ± 0.072 | -0.428 | 0.669 |
| Generation length (residual) | 0.051 ± 0.029 | 1.768 | 0.077 |
| Migration (migratory vs sedentary) | 0.017 ± 0.023 | 0.749 | 0.454 |
| | | lambda = 0.963; r2 = 0.135 | |
| **Model: mPC2 ~ other traits** | | | |
| (Intercept) | -0.205 ± 0.247 | -0.829 | 0.407 |
| Body mass (log-10) | 0.021 ± 0.043 | 0.488 | 0.626 |
| **Brain size (residual)** | 0.274 ± 0.040 | 6.801 | <0.001 |
| **Eye size (residual)** | 0.575 ± 0.289 | 1.990 | 0.047 |
| **ePC1 (Arboreal frugivory)** | 0.118 ± 0.023 | 5.105 | <0.001 |
| ePC2 (Aquatic predation) | -0.041 ± 0.031 | -1.316 | 0.189 |
| **ePC3 (Land Predation)** | -0.066 ± 0.031 | -2.145 | 0.032 |
| **ePC4 (Insectivory)** | -0.071 ± 0.029 | -2.435 | 0.015 |
| **sPC1 (Bonding)** | 0.077 ± 0.024 | 3.266 | 0.001 |
| sPC2 (Grouping) | 0.030 ± 0.020 | 1.480 | 0.139 |
| **Parental providing (log +1)** | 0.198 ± 0.060 | 3.309 | <0.001 |
| Generation length (residual) | -0.004 ± 0.024 | -0.149 | 0.882 |
| **Migration (migratory vs sedentary)** | 0.039 ± 0.019 | 2.081 | 0.038 |
| | | lambda = 0.952; r2 = 0.181 | |
| **Model: mPC3 ~ other traits** | | | |
| (Intercept) | -0.271 ± 0.299 | -0.908 | 0.364 |
| **Body mass (log-10)** | 0.222 ± 0.047 | 4.755 | <0.001 |
| **Brain size (residual)** | 0.189 ± 0.043 | 4.392 | <0.001 |
| Eye size (residual) | 0.229 ± 0.301 | 0.759 | 0.448 |
| ePC1 (Arboreal frugivory) | -0.009 ± 0.025 | -0.360 | 0.719 |
| ePC2 (Aquatic predation) | 0.057 ± 0.034 | 1.695 | 0.091 |
| **ePC3 (Land Predation)** | 0.077 ± 0.033 | 2.322 | 0.021 |

| | | | |
|---|---|---|---|
| ePC4 (Insectivory) | 0.058 ± 0.031 | 1.906 | 0.057 |
| sPC1 (Bonding) | -0.014 ± 0.026 | -0.549 | 0.583 |
| sPC2 (Grouping) | -0.028 ± 0.021 | -1.339 | 0.181 |
| Parental providing (log +1) | -0.081 ± 0.062 | -1.303 | 0.193 |
| Generation length (residual) | -0.003 ± 0.026 | -0.098 | 0.922 |
| **Migration (migratory vs sedentary)** | -0.064 ± 0.020 | -3.296 | 0.001 |
| | | lambda = 0.982; r2 = 0.118 | |
| **Model: ePC1 ~ other traits** | | | |
| (Intercept) | 0.116 ± 0.429 | 0.270 | 0.787 |
| Body mass (log-10) | -0.016 ± 0.076 | -0.210 | 0.834 |
| Brain size (residual) | 0.008 ± 0.075 | 0.110 | 0.912 |
| Eye size (residual) | 0.908 ± 0.523 | 1.734 | 0.083 |
| mPC1 (Locomotion) | -0.009 ± 0.058 | -0.162 | 0.871 |
| **mPC2 (Beak thickness)** | 0.308 ± 0.070 | 4.413 | <0.001 |
| mPC3 (Long beak & leg) | 0.009 ± 0.062 | 0.147 | 0.883 |
| **sPC1 (Bonding)** | -0.121 ± 0.042 | -2.879 | 0.004 |
| sPC2 (Grouping) | -0.004 ± 0.036 | -0.112 | 0.911 |
| **Parental providing (log +1)** | 0.264 ± 0.107 | 2.469 | 0.014 |
| Generation length (residual) | 0.034 ± 0.043 | 0.794 | 0.428 |
| Migration (migratory vs sedentary) | -0.039 ± 0.034 | -1.128 | 0.260 |
| | | lambda = 0.942; r2 = 0.070 | |
| **Model: ePC2 ~ other traits** | | | |
| (Intercept) | 0.193 ± 0.332 | 0.581 | 0.562 |
| **Body mass (log-10)** | 0.149 ± 0.057 | 2.633 | 0.009 |
| Brain size (residual) | -0.028 ± 0.056 | -0.510 | 0.610 |
| Eye size (residual) | 0.153 ± 0.384 | 0.399 | 0.690 |
| **mPC1 (Locomotion)** | -0.149 ± 0.043 | -3.460 | <0.001 |
| mPC2 (Beak thickness) | -0.034 ± 0.052 | -0.650 | 0.516 |
| **mPC3 (Long beak & leg)** | 0.092 ± 0.047 | 1.971 | 0.049 |
| sPC1 (Bonding) | -0.008 ± 0.031 | -0.260 | 0.795 |
| sPC2 (Grouping) | 0.020 ± 0.026 | 0.766 | 0.444 |
| Parental providing (log +1) | 0.069 ± 0.079 | 0.877 | 0.381 |
| **Generation length (residual)** | 0.062 ± 0.031 | 1.964 | 0.050 |
| Migration (migratory vs sedentary) | 0.001 ± 0.025 | 0.048 | 0.962 |
| | | lambda = 0.957; r2 = 0.039 | |
| **Model: ePC3 ~ other traits** | | | |
| (Intercept) | -0.187 ± 0.289 | -0.647 | 0.518 |
| **Body mass (log-10)** | 0.174 ± 0.054 | 3.225 | 0.001 |

| | | | |
|---|---|---|---|
| Brain size (residual) | -0.014 ± 0.055 | -0.256 | 0.798 |
| **Eye size (residual)** | 1.206 ± 0.386 | 3.123 | 0.002 |
| mPC1 (Locomotion) | 0.055 ± 0.042 | 1.318 | 0.188 |
| mPC2 (Beak thickness) | -0.069 ± 0.050 | -1.366 | 0.172 |
| mPC3 (Long beak & leg) | 0.082 ± 0.044 | 1.848 | 0.065 |
| sPC1 (Bonding) | -0.008 ± 0.030 | -0.276 | 0.782 |
| **sPC2 (Grouping)** | -0.073 ± 0.026 | -2.745 | 0.006 |
| Parental providing (log +1) | 0.119 ± 0.078 | 1.531 | 0.126 |
| Generation length (residual) | -0.029 ± 0.031 | -0.916 | 0.360 |
| Migration (migratory vs sedentary) | 0.027 ± 0.025 | 1.071 | 0.284 |
| | | lambda = 0.912; r2 = 0.077 | |
| **Model: ePC4 ~ other traits** | | | |
| (Intercept) | 0.101 ± 0.291 | 0.349 | 0.727 |
| **Body mass (log-10)** | -0.408 ± 0.056 | -7.240 | <0.001 |
| Brain size (residual) | 0.002 ± 0.058 | 0.042 | 0.967 |
| **Eye size (residual)** | 0.942 ± 0.413 | 2.280 | 0.023 |
| mPC1 (Locomotion) | -0.009 ± 0.044 | -0.211 | 0.833 |
| mPC2 (Beak thickness) | -0.101 ± 0.053 | -1.904 | 0.057 |
| **mPC3 (Long beak & leg)** | 0.111 ± 0.046 | 2.383 | 0.017 |
| sPC1 (Bonding) | -0.036 ± 0.032 | -1.126 | 0.261 |
| sPC2 (Grouping) | -0.042 ± 0.028 | -1.498 | 0.135 |
| Parental providing (log +1) | 0.147 ± 0.082 | 1.792 | 0.074 |
| Generation length (residual) | 0.034 ± 0.033 | 1.029 | 0.304 |
| Migration (migratory vs sedentary) | 0.038 ± 0.027 | 1.390 | 0.165 |
| | | lambda = 0.887; r2 = 0.105 | |
| **Model: sPC1 ~ other traits** | | | |
| (Intercept) | 0.043 ± 0.412 | 0.103 | 0.918 |
| **Body mass (log-10)** | -0.240 ± 0.073 | -3.286 | 0.001 |
| Brain size (residual) | 0.087 ± 0.069 | 1.260 | 0.208 |
| Eye size (residual) | -0.045 ± 0.485 | -0.092 | 0.927 |
| mPC1 (Locomotion) | -0.035 ± 0.054 | -0.646 | 0.518 |
| **mPC2 (Beak thickness)** | 0.201 ± 0.066 | 3.051 | 0.002 |
| mPC3 (Long beak & leg) | -0.039 ± 0.059 | -0.671 | 0.503 |
| **ePC1 (Arboreal frugivory)** | -0.106 ± 0.039 | -2.746 | 0.006 |
| ePC2 (Aquatic predation) | 0.032 ± 0.052 | 0.614 | 0.539 |
| ePC3 (Land Predation) | -0.016 ± 0.051 | -0.316 | 0.752 |
| ePC4 (Insectivory) | -0.033 ± 0.048 | -0.694 | 0.488 |
| **Parental providing (log +1)** | 0.272 ± 0.099 | 2.757 | 0.006 |

| | | | |
|---|---|---|---|
| Generation length (residual) | 0.073 ± 0.039 | 1.854 | 0.064 |
| **Migration (migratory vs sedentary)** | -0.063 ± 0.031 | -2.017 | 0.044 |
| | | lambda = 0.954; r2 = 0.076 | |
| **Model: sPC2 ~ other traits** | | | |
| (Intercept) | -0.240 ± 0.289 | -0.830 | 0.407 |
| **Body mass (log-10)** | 0.173 ± 0.074 | 2.342 | 0.019 |
| Brain size (residual) | -0.099 ± 0.077 | -1.282 | 0.200 |
| Eye size (residual) | -1.092 ± 0.584 | -1.868 | 0.062 |
| mPC1 (Locomotion) | 0.041 ± 0.057 | 0.734 | 0.463 |
| mPC2 (Beak thickness) | 0.121 ± 0.069 | 1.751 | 0.080 |
| **mPC3 (Long beak & leg)** | -0.135 ± 0.057 | -2.351 | 0.019 |
| ePC1 (Arboreal frugivory) | -0.020 ± 0.042 | -0.490 | 0.624 |
| ePC2 (Aquatic predation) | 0.091 ± 0.053 | 1.714 | 0.087 |
| **ePC3 (Land Predation)** | -0.157 ± 0.055 | -2.830 | 0.005 |
| ePC4 (Insectivory) | -0.079 ± 0.053 | -1.491 | 0.137 |
| Parental providing (log +1) | -0.072 ± 0.102 | -0.710 | 0.478 |
| Generation length (residual) | 0.055 ± 0.045 | 1.237 | 0.217 |
| Migration (migratory vs sedentary) | 0.007 ± 0.038 | 0.178 | 0.859 |
| | | lambda = 0.698; r2 = 0.060 | |
| **Model: Time fed ~ other traits** | | | |
| (Intercept) | -0.355 ± 0.148 | -2.401 | 0.017 |
| **Body mass (log-10)** | 0.150 ± 0.028 | 5.377 | <0.001 |
| **Brain size (residual)** | 0.070 ± 0.027 | 2.557 | 0.011 |
| Eye size (residual) | -0.087 ± 0.195 | -0.446 | 0.656 |
| mPC1 (Locomotion) | 0.010 ± 0.021 | 0.458 | 0.647 |
| **mPC2 (Beak thickness)** | 0.090 ± 0.026 | 3.496 | <0.001 |
| mPC3 (Long beak & leg) | -0.040 ± 0.023 | -1.761 | 0.079 |
| ePC1 (Arboreal frugivory) | 0.028 ± 0.015 | 1.865 | 0.063 |
| ePC2 (Aquatic predation) | 0.012 ± 0.020 | 0.602 | 0.547 |
| ePC3 (Land Predation) | 0.035 ± 0.020 | 1.734 | 0.083 |
| ePC4 (Insectivory) | 0.026 ± 0.019 | 1.382 | 0.167 |
| **sPC1 (Bonding)** | 0.043 ± 0.015 | 2.830 | 0.005 |
| sPC2 (Grouping) | -0.013 ± 0.013 | -1.010 | 0.313 |
| **Generation length (residual)** | 0.034 ± 0.016 | 2.168 | 0.030 |
| **Migration (migratory vs sedentary)** | -0.043 ± 0.013 | -3.446 | <0.001 |
| | | lambda = 0.922; r2 = 0.170 | |
| **Model: Generation length ~ other traits** | | | |
| (Intercept) | 0.027 ± 0.328 | 0.082 | 0.935 |

| | | | |
|---|---|---|---|
| Body mass (log-10) | 0.000 ± 0.070 | 0.000 | 1.000 |
| **Brain size (residual)** | 0.202 ± 0.069 | 2.948 | 0.003 |
| Eye size (residual) | 0.000 ± 0.501 | 0.000 | 1.000 |
| **mPC1 (Locomotion)** | 0.106 ± 0.052 | 2.034 | 0.042 |
| mPC2 (Beak thickness) | -0.019 ± 0.064 | -0.300 | 0.764 |
| mPC3 (Long beak & leg) | 0.000 ± 0.055 | 0.003 | 0.998 |
| ePC1 (Arboreal frugivory) | 0.007 ± 0.038 | 0.181 | 0.856 |
| ePC2 (Aquatic predation) | 0.080 ± 0.049 | 1.634 | 0.103 |
| ePC3 (Land Predation) | -0.039 ± 0.050 | -0.779 | 0.436 |
| ePC4 (Insectivory) | 0.037 ± 0.048 | 0.766 | 0.444 |
| sPC1 (Bonding) | 0.065 ± 0.038 | 1.696 | 0.090 |
| sPC2 (Grouping) | 0.032 ± 0.034 | 0.954 | 0.340 |
| **Parental providing (log +1)** | 0.241 ± 0.097 | 2.495 | 0.013 |
| Migration (migratory vs sedentary) | -0.030 ± 0.032 | -0.927 | 0.354 |
| | | lambda = 0.863; r2 = 0.063 | |

**Table S8.** Variance inflation factor (VIF) analyses (using the function *vif.phylolm*, see https://github.com/mrhelmus/phylogeny_manipulation) of all parameters included in the combined model.

| **Parameter** | **VIF for combined model in Table S3** | **VIF for combined model in Table S4** |
|---|---|---|
| Body mass (log-10) | 1.247 | 1.368 |
| Eye size (residual) | | 1.117 |
| mPC1 (Locomotion) | 1.200 | 1.230 |
| mPC2 (Beak thickness) | 1.192 | 1.205 |
| mPC3 (Long beak & leg) | 1.156 | 1.133 |
| Parental providing (log +1) | 1.204 | 1.204 |
| Generation length (residual) | 1.056 | 1.051 |
| Migration (migratory vs sedentary) | 1.078 | 1.084 |
| ePC1 (Arboreal frugivory) | 1.215 | 1.218 |
| ePC2 (Aquatic predation) | 1.248 | 1.166 |
| ePC3 (Land Predation) | 1.060 | 1.123 |
| ePC4 (Insectivory) | 1.104 | 1.181 |
| sPC1 (Bonding) | 1.061 | 1.084 |
| sPC2 (Grouping) | 1.039 | 1.048 |

**Table S9**. Variable-rates model with uniform prior distributions and the iterations for single-chain Markov chain Monte Carlo adjusted to variables. We assessed the convergence of the chains using the Cramer-von-Mises statistic implemented in the R package coda[4]. Both the stationarity test and the half-width test were applied to evaluate the convergence of the chain's likelihood, the phylogenetic mean, the Brownian motion variance, the number of branch lengths scaled using variable rates and the number using variable rates of nodes scaled using variable rates. We note that all chains for each trait passed these tests, except for the parental provisioning, where the phylogenetic mean (Alpha) and Brownian motion variance (Sigma^2) did not pass the half-width test. Nonetheless, the simulation of the variable-rates model for parental provision is the best one (see Table S9(, and thus, we used this model to reconstruct the ancestral state for parental provision.

| Trait | Prior 1 | Prior 2 | Iterations | Brun in | Sample | Stationarity | Halfwidth |
|---|---|---|---|---|---|---|---|
| Brain size | Alpha-1 Uniform -1 1 | Sigma-1 Uniform 0 0.001 | $10 \times 10^8$ | $10^7$ | $10^5$ | All passed | All passed |
| mPC1 | Alpha-1 Uniform -1 1 | Sigma-1 Uniform 0 0.001 | $10 \times 10^8$ | $10^7$ | $10^5$ | All passed | All passed |
| mPC2 | Alpha-1 Uniform -1 1 | Sigma-1 Uniform 0 0.001 | $10 \times 10^8$ | $10^7$ | $10^5$ | All passed | All passed |
| mPC3 | Alpha-1 Uniform -3 4 | Sigma-1 Uniform 0 0.001 | $15 \times 10^8$ | $10^7$ | $10^5$ | All passed | All passed |
| Parental providing | Alpha-1 Uniform 0 6 | Sigma-1 Uniform 0 2 | $10 \times 10^8$ | $10^7$ | $10^5$ | All passed | Alpha, Sigma^2 |
| Generation length | Alpha-1 Uniform -0.6 0.6 | Sigma-1 Uniform 0 0.001 | $10 \times 10^8$ | $10^7$ | $10^5$ | All passed | All passed |
| Migration | q01 uniform 0 10 | q10 uniform 0 10 | $5 \times 10^8$ | $5 \times 10^6$ | $10^4$ | All passed | All passed |

**Table S10**. Comparison of model fits for different models of traits evolution and phylogenetic signal for each trait complex using the time-calibrated species tree. The performance of each model was quantified via two metrics: log-likelihood difference presented outside brackets, i.e., the difference between the highest log-likelihood among the five models analysed, and the model's log-likelihood via the AIC difference shown in brackets, i.e., the difference between the model's AIC value and the lowest AIC value across all models. We note that Variable rates has highest log-likelihood and lowest AIC for all traits, hence its value is 0(0).

| | Model comparison log-likelihood difference (AIC difference) | | | | | | |
|---|---|---|---|---|---|---|---|
| **Trait** | **White noise** | **Brownian motion** | **Omstein-Uhlenbeck** | **Early burst** | **Variable rates** | ***λ*** | ***K*** |
| Brain size | 979 (1646) | 421 (528) | 342 (374) | 421 (530) | 0 (0) | 0.913 | 0.49 |
| mPC1 | 1113 (1910) | 425 (535) | 391 (469) | 425 (537) | 0 (0) | 0.972 | 0.467 |
| mPC2 | 1021 (1732) | 465 (620) | 408 (507) | 465 (622) | 0 (0) | 0.958 | 0.437 |
| mPC3 | 1036 (1770) | 462 (622) | 407 (515) | 462 (624) | 0 (0) | 0.959 | 0.381 |
| Parental providing | 13779 (26176) | 12288 (23194) | 12286 (23191) | 12288 (23196) | 0 (0) | 0.96 | 1.857 |
| Generation length | 713 (1098) | 807 (1287) | 536 (746) | 807 (1289) | 0 (0) | 0.876 | 0.121 |
| Migration | 247 (194) | 513 (726) | 199 (98) | 513 (728) | 0 (0) | 0.657 | 0.0 |

**Table S11**. Effective sample sizes of number of shift events and log likelihoods from BAMM analysis checked via the R package coda[4].

| | No. of events | Log likelihoods |
|---|---|---|
| **Analysis with 2133 species** | | |
| Brain size | 2031.27 | 1329.65 |
| mPC1 | 1541.64 | 1213.67 |
| mPC2 | 1678.20 | 1371.54 |
| mPC3 | 1437.48 | 1239.21 |
| **Analysis with 1199 species** | | |
| Brain size | 2293.84 | 2153.39 |
| Eye size | 3341.84 | 2482.57 |
| mPC1 | 3566.73 | 2724.14 |
| mPC2 | 3560.43 | 2265.15 |
| mPC3 | 1789.24 | 1282.66 |

**Figures**

**Figure S1**. Phylogenetic associations among morphology, eye size, life-history traits, eco-social niche and brain size (n = 660). a, presents a heatmap matrix illustrating the relationships between various traits. For each trait in the row, a model was constructed against the other 14 column variables, resulting in a total of 13 models. All models accounted for phylogenetic relationships and included body mass (log-10) as a consistent fixed variable. The models were developed using Phylogenetic Generalized Least Squares (PGLS) via the 'phylolm' package[3]. Within each matrix cell, t-values are displayed and their magnitude is visually represented by the colour intensity of the cell. Cells with black-coloured t-values indicate statistical significance after Bonferroni correction, corresponding to a p-value threshold of less than 0.05/13. b, significant Bonferroni-corrected pair-wise associations among morphology including eye size (yellow), life history (blue), ecology and sociality (green) and relative brain size (orange); solid lines represent relationships that are significant in both directions, broken lines represent relationships significant only in one direction (migration could only be included as an independent variable).

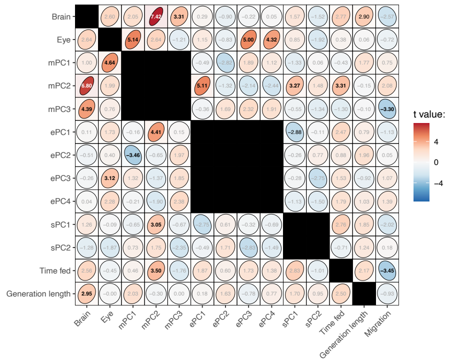

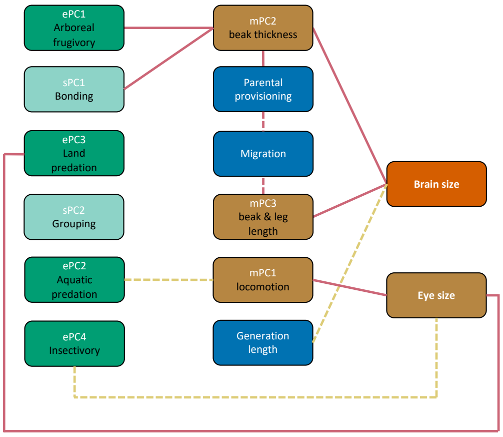

**Figure S2.** Estimated coefficients and effect sizes of phylogenetically controlled mixed models of the morphology model, the eco-social model, life-history model, and the combined model on relative brain size in birds. Colour-filled circles denote estimated effects, lines denote the 95% lower and upper confidence limits generated in the R package MCMCglmm[2] based on a consensus phylogeny (see methods for more details). The corresponding full models are shown in Table S4.

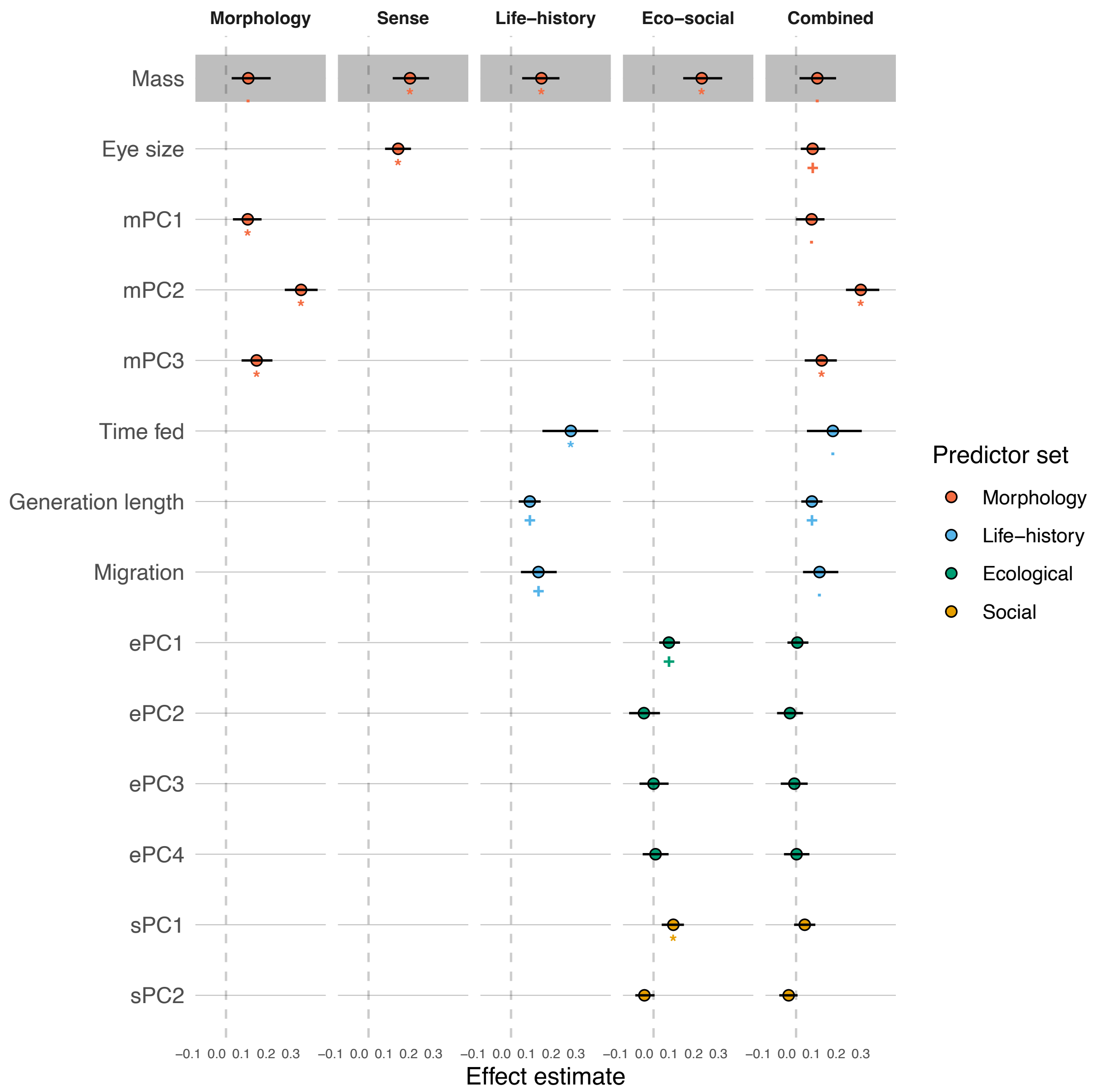

**Figure S3**. a) median rates and rate distributions for net phenotypic (brain size (residual), eye size(residual) and mPC1-3) macroevolutionary rate. the y axis is scaled from zero to the maximum median rate for all dataset, for relative comparability. b) Rates of phenotypic evolution in bird, branches are coloured in a rainbow scale by Jenks natural breaks method. Red dots at nodes represent diversification shifts in the maximum credibility set.

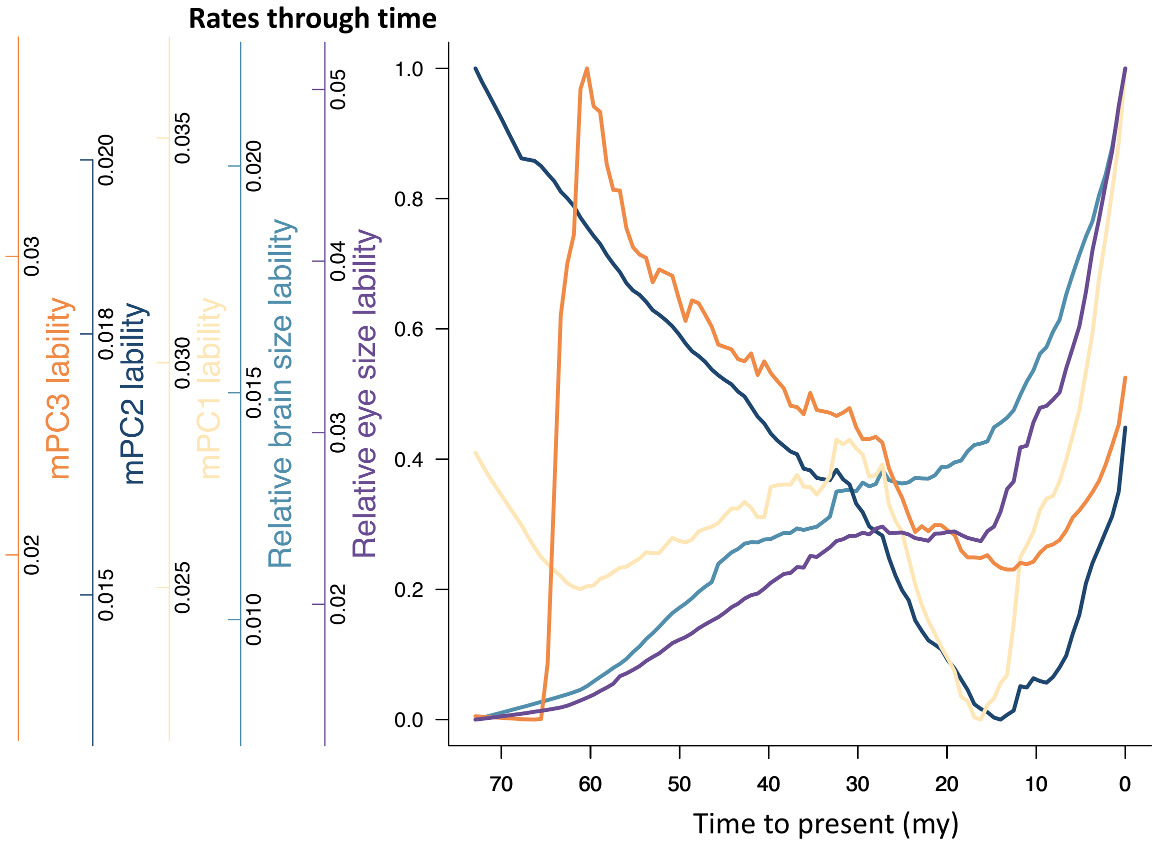

Eye

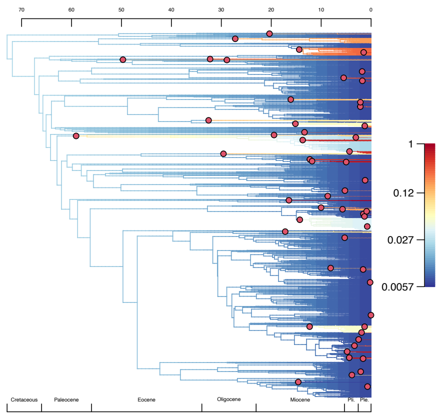

Brain

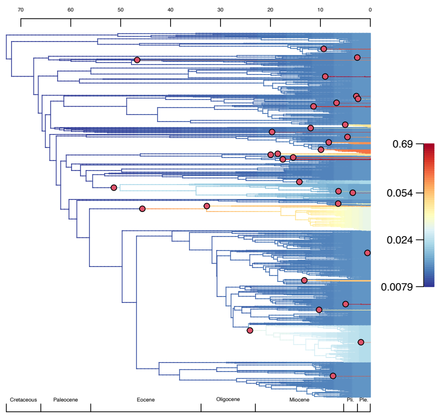

mPC1

mPC2

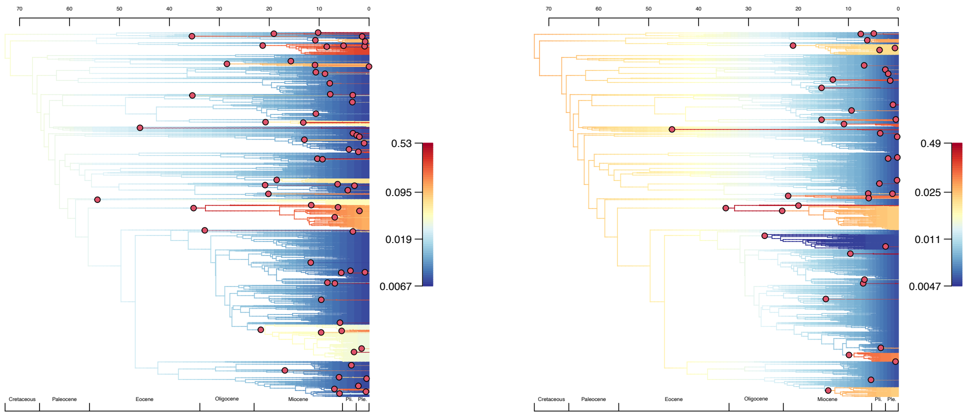

mPC3

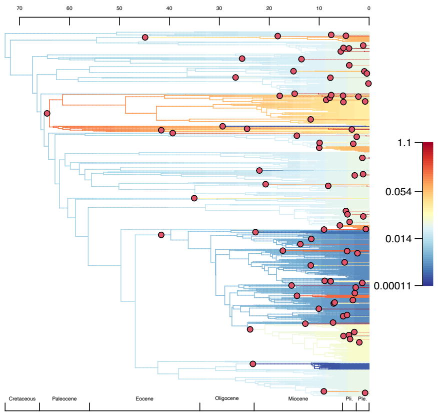

### References

1. Revelle, W. psych: Procedures for personality and psychological research. Preprint at (2017).
2. Hadfield, J. D. MCMC methods for multi-response generalized linear mixed models: the MCMCglmm R package. *J Stat Softw* **33**, 1–22 (2010).
3. Ho, L. si T. & Ane, C. A Linear-Time Algorithm for Gaussian and Non-Gaussian Trait Evolution Models. *Syst Biol* **63**, 397–408 (2014).
4. Plummer, M., Best, N., Cowles, K. & Vines, K. CODA: convergence diagnosis and output analysis for MCMC. *R New* **6**, 7–11 (2005).
